# Two brain pathways initiate distinct forward walking programs in *Drosophila*

**DOI:** 10.1101/798439

**Authors:** Salil S. Bidaye, Meghan Laturney, Amy K. Chang, Yuejiang Liu, Till Bockemühl, Ansgar Büschges, Kristin Scott

## Abstract

An animal at rest or engaged in stationary behaviors can instantaneously initiate goal-directed walking. How descending brain inputs trigger rapid transitions from a non-walking state to an appropriate walking state is unclear. Here, we identify two specific neuronal classes in the *Drosophila* brain that drive two distinct forward walking programs in a context-specific manner. The first class, named P9, consists of descending neurons that drive forward walking with ipsilateral turning. P9 receives inputs from central courtship-promoting neurons and visual projection neurons and is necessary for a male to track a female during courtship. The second class comprises novel, higher order neurons, named BPN, that drives straight, forward walking. BPN is required for high velocity walking and is active during long, fast, straight walking bouts. Thus, this study reveals separate brain pathways for object-directed steering and fast straight walking, providing insight into how the brain initiates different walking programs.

## Introduction

Animals show a remarkable ability to seamlessly adjust their walking patterns in response to dynamic sensory stimuli, as they move about an unpredictable environment. Moreover, animals not only are capable of modifying an ongoing walking pattern, but also have the ability to transition from a non-walking state directly into a goal-appropriate walking state. For example, an animal may suddenly spring into an escape run at the sight of a predator or initiate a spontaneous exploratory walking bout. However, it is unclear how the nervous system instantaneously generates such a complex and varied locomotor output.

In both vertebrates and invertebrates, the leg movements that comprise a walking pattern are hypothesized to be controlled by a distributed network of pattern generating modules in the spinal cord or nerve cord, with each module controlling the movement of each leg joint (Brown, 1911; Büschges et al., 1995; Cheng et al., 1998; Grillner and Zangger, 1975; Hägglund et al., 2013; Ryckebusch and Laurent, 1993). Sensory feedback as well as central neural circuits are crucial for coordinating this network in order to generate walking (Bässler and Büschges, 1998; Shik and Orlovsky, 1976; Tuthill and Azim, 2018). These leg movement control circuits exist in a fundamentally different functional state in a non-walking animal versus a walking animal. For example, classical studies show that the same force applied to the leg of a non-moving animal produces a distinct motor output compared to that of an animal producing voluntary leg movements (Bässler, 1976; Pearson, 1995). Similarly, inter-leg coordination during stationary behaviors like grooming (Mueller et al., 2019) or searching (Berg et al., 2015) is completely different from that during walking. Despite these distinct functional states, descending inputs from the brain are able to instantaneously switch the spinal cord or nerve cord circuits from a non-walking state into a walking state (Bidaye et al., 2018; Jordan et al., 2008; Ruder and Arber, 2019). Whether walking is initiated by concerted population activity of descending inputs that coordinate pattern generating modules in the spinal cord or whether a single brain command is able to drive such a complex state change is unknown. Furthermore, it is unclear whether walking initiated in different contexts, such as escaping, pursuing or exploring, results from distinct descending inputs or from modulation of a common descending pathway for walking initiation. To distinguish potential brain mechanisms for locomotor control and the generation of context-dependent behavior, it is essential to elucidate descending pathways that encode walking initiation.

In mammals, brainstem circuits are critical for walking initiation. The mesencephalic locomotor region (MLR) is one such brainstem region implicated in walking initiation across different mammalian species (Shik and Orlovsky, 1976; Skinner and Garcia-Rill, 1984). The MLR activates neurons in the reticular formation in the hindbrain that in turn drive spinal cord networks for locomotion. Recent studies have begun delineating mouse MLR and brainstem regions into molecularly defined subsets that control specific aspects of locomotion (Bouvier et al., 2015; Caggiano et al., 2018; Capelli et al., 2017). However, given the heterogeneity of these brain areas in terms of functions, cell-types, and projection targets, it remains challenging to examine how descending pathways recruit spinal cord circuits for walking initiation.

In insects, the small number of ~300-500 descending neurons (DNs) that project to the ventral nerve cord (VNC) provides the opportunity to finely probe how locomotor control is encoded in descending pathways. Although specific DNs have been associated with specific aspects of walking including walking initiation in different insect species (Ache et al., 2015; Böhm and Schildberger, 1992; Burdohan and Comer, 1996; Zorović and Hedwig, 2013), lack of reproducible access to these neurons makes it difficult to examine their function. In addition, little is known about how higher brain neurons control context-specific walking initiation in insects. The sophisticated genetic tools in *Drosophila melanogaster* allow reproducible access to specific neurons including DNs (Namiki et al., 2018) and greatly facilitate functional characterization. Recent work revealed that many DNs are able to modify an ongoing walking program (Bidaye et al., 2014; Cande et al., 2018). In addition, the activity of a few DNs has been shown to correlate with locomotor patterns (Chen et al., 2018; Tschida and Bhandawat, 2015). However, it is still unclear if specific brain inputs, both at the level of DNs as well as higher brain areas, are capable of initiating a coordinated walking pattern and how distinct walking initiations are manifested in different behavioral contexts.

Here, we leveraged recently developed genetic tools and coupled them with novel behavioral assays and functional imaging to examine walking initiation neurons in *Drosophila*. From a targeted optogenetic screen for walking initiation neurons, we identified two neuronal types that initiate two distinct modes of forward walking: P9 DN induces forward walking with an ipsilateral turning component, whereas novel higher brain neurons, which we name Bolt protocerebral neurons (BPN), induce straight forward walking. Functional connectivity and neural silencing studies revealed that the ipsilateral turning walking program driven by P9 is essential for object directed walking in the context of courtship. In contrast, BPNs were dispensable for courtship. Instead, *in-vivo* imaging and behavioral experiments showed that BPNs are specifically necessary for high velocity, straight forward walking. These studies show that the activity of specific brain neurons is sufficient to switch downstream motor control circuits from a non-walking state into a walking state and provide evidence for distinct descending pathways for walking initiations in different contexts.

## Results

### A neural activation screen identifies candidate walking initiation neurons

To examine how animals initiate walking rather than modify an ongoing walking pattern, we designed an optogenetics based assay to screen for neurons that trigger walking in non-walking flies. Flies spend most of their time walking when restricted to a small arena where they cannot fly. To bias flies toward a non-walking state, flies were dusted with fine powder to induce grooming (Seeds et al., 2014). This greatly reduced baseline walking activity, an essential condition to screen for walking initiation.

Descending neurons (DNs) send information from the brain to the VNC and represent a critical relay for motor control. We therefore screened 156 lines that targeted potential DNs, consisting of the DN split-Gal4 collection which covers over a third of all Drosophila DNs, (Namiki et al., 2018) as well as selected Gal4 lines from the Janelia and VT collections that label DNs (Bidaye et al., 2014; Jenett et al., 2012; Tirian and Dickson, 2017). Powdered and non-powdered flies expressing the red-shifted channelrhodopsin CsChrimson (Klapoetke et al., 2014) in neural candidates were stimulated with light and locomotion of single flies was monitored (Figure 1A). Three lines increased walking more than 5 fold relative to controls when covered with powder and more than 1.2 fold when not powdered (Figures 1B and S1A). Upon retest with acute stimulation, two of the lines, SS01540 and SS01587 from the DN split-Gal4 collection (Namiki et al., 2018), reliably initiated walking in powdered flies (Figures 1C, S1B and Video S1).

**Figure 1:**
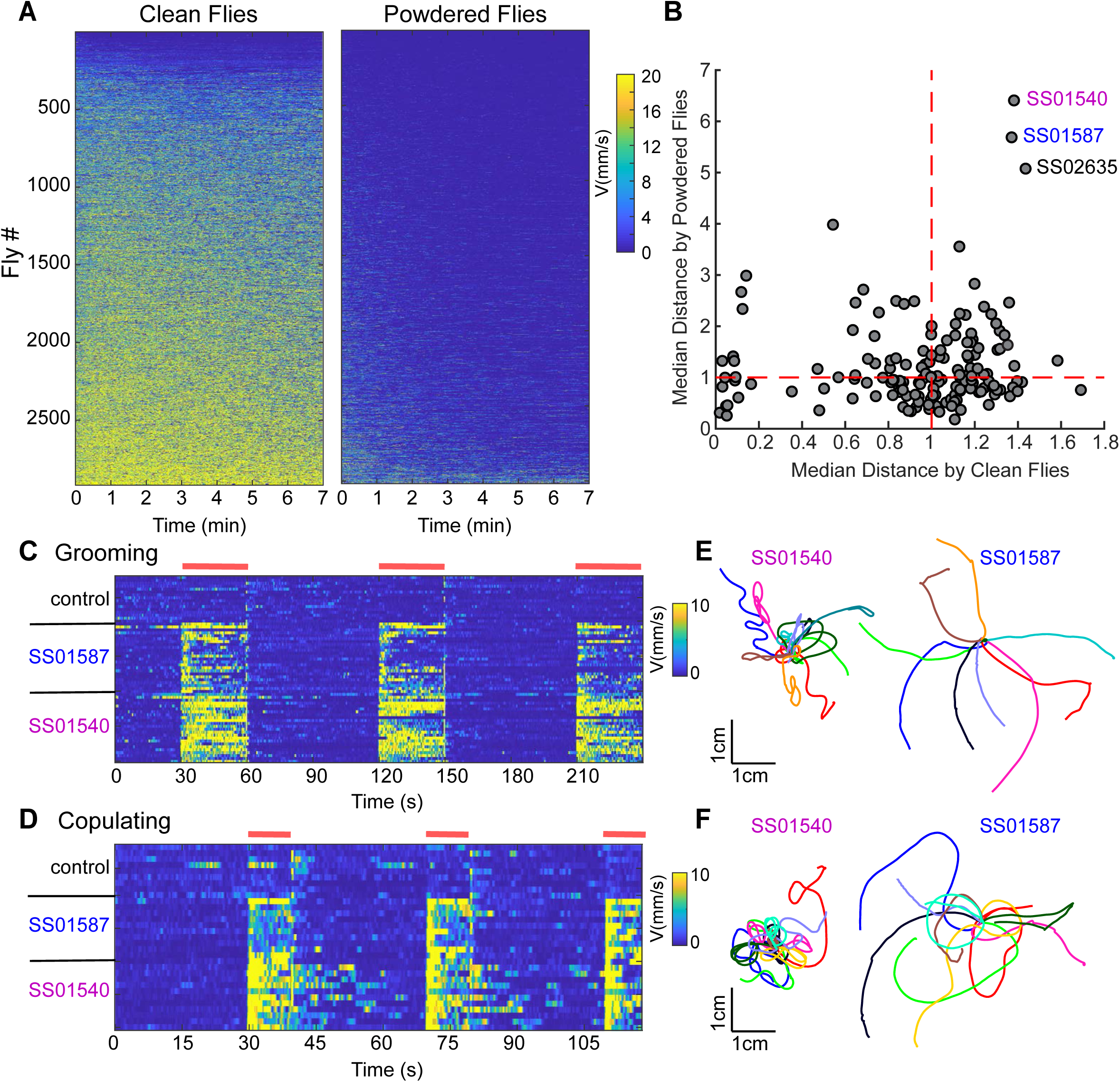
Optogenetic screen identifies candidate walking initiation lines. (A) Translational velocity of non-powdered, “clean” flies (left) and “powdered”, grooming flies (right) upon optogenetic activation with CsChrimson during 7 minute assay, shown as velocity heatmaps for individual flies arranged from lowest to highest mean velocity. (B) Normalized median distance traveled by clean versus powdered flies for each genotype. n=12-16 per genotype per condition. (C) Walking initiation upon transient activation in grooming flies for control, SS01540 and SS01587, shown as velocity heatmaps for individual flies (red bars indicate light ON). n=16-24 flies/genotype. (D) Walking initiation of females upon transient activation in copulating flies, shown as velocity heatmaps for individual flies (red bars indicate light ON). n=9-11 flies/genotype. (E) Example walking trajectories, 4 seconds lightON, for SS01540 and SS01587 in the grooming assay, 10 flies/genotype. (F) Example walking trajectories, 4 seconds lightON, for SS01540 and SS01587 in the copulation assay, 10 flies/genotype. See Figure S1 for statistical analysis of candidate lines.

If neurons in these lines are bona fide walking initiation neurons, then neural activation should lead to walking in a context other than grooming. Copulation is a persistent non-walking state in which the male and the female remain stationary for extended periods (Crickmore and Vosshall, 2013), although the female posture is compatible with walking. We therefore expressed CsChrimson in SS01540 and SS01587 and transiently activated the neurons of females mid-copulation. Remarkably, the females reliably initiated walking while remaining conjoined with the mounted male (Figures 1D, S1C and Video S2). This demonstrates that activation of specific neurons in the fly initiates walking and overrides competing behavioral states, including grooming and copulating.

Although both lines induced forward walking, SS01540 and SS01587 induced forward walks with different trajectories (Figures 1E, 1F and Video S1). SS01540 activation elicited forward walking with a strong turning component whereas SS01587 activation induced straight forward walking, for both grooming and copulation assays (Figures 1E, 1F, S1D, and S1E). Thus, via a targeted optogenetic screen, we identified two walking initiation candidates that induce two modes of forward walking, suggesting that they might represent activation of distinct walking initiation pathways.

### P9 DNs initiate forward walking with an ipsilateral turning component

To begin to examine neural pathways that initiate forward walking, we focused on SS01540 as its expression pattern is sparse. This line labels DNp09 (henceforth referred to as P9) (Namiki et al., 2018), a pair of ipsilateral DNs (one cell per hemisphere) and a few additional neurons (Figures S2A and S2B). We generated a second P9 split-Gal4 line (P9S2) that targets P9 but does not share off-target cells with SS01540 (Figures S2C and S2D). Activating P9S2 neurons induced a walking initiation phenotype with a strong turning component similar to SS01540 (Figure S2E), confirming the role of P9 in walking initiation.

SS01540 activation has previously been reported to induce transient walking and freezing/stopping behaviors (Cande et al., 2018; Zacarias et al., 2018). We reproduced the phenotype that SS01540>CsChrimson(attp2) flies showed an initial increase in walking velocity followed by freezing upon activation (Figure S2F), using published methods. However, freezing was not observed in three other genetic driver and effector combinations, although all genotypes strongly label P9 and initiate walking on activation (Figures S2A-D and S2F). Whether the freezing phenotype of SS01540>CsChrimson(attp2) is a consequence of high CsChrimson expression in P9 or off-target neurons (Figure S2G) is unclear. Regardless, in all cases, optogenetic stimulation increased both forward and angular velocity, corroborating that P9 activation elicits walking.

P9 dendrites are in the posterior protocerebrum and presynaptic outputs in the subesophageal zone (SEZ) and the ipsilateral leg VNC neuropils (Figures 2A and 2B). The ipsilateral dendrites and axons suggest that activation of a single P9 neuron might lead to turning in a specific direction. To test this, mosaic animals with CsChrimson stochastically expressed in one or both P9 neurons were generated, dusted with powder, and stimulated with red light while monitoring locomotion. Several flies turned only to the left or right upon activation (Video S3), whereas other flies turned in both directions. All animals turning in both directions had bilateral P9 expression (Figure 2C), whereas all animals that turned right expressed CsChrimson unilaterally in the right P9 (Figure 2D) and vice versa for left turning individuals (Figure 2E). The stochastic labeling and manipulation of single P9 neurons demonstrates that P9 is the causal neuron in the split-Gal4 line that initiates a complex locomotor program of forward walking with ipsilateral turns. Moreover, these studies reveal that unilateral activation of a single P9 neuron initiates ipsilateral forward turns and raise the possibility that P9 activity may be recruited to initiate walking towards an ipsilateral attractive stimulus.

**Figure 2:**
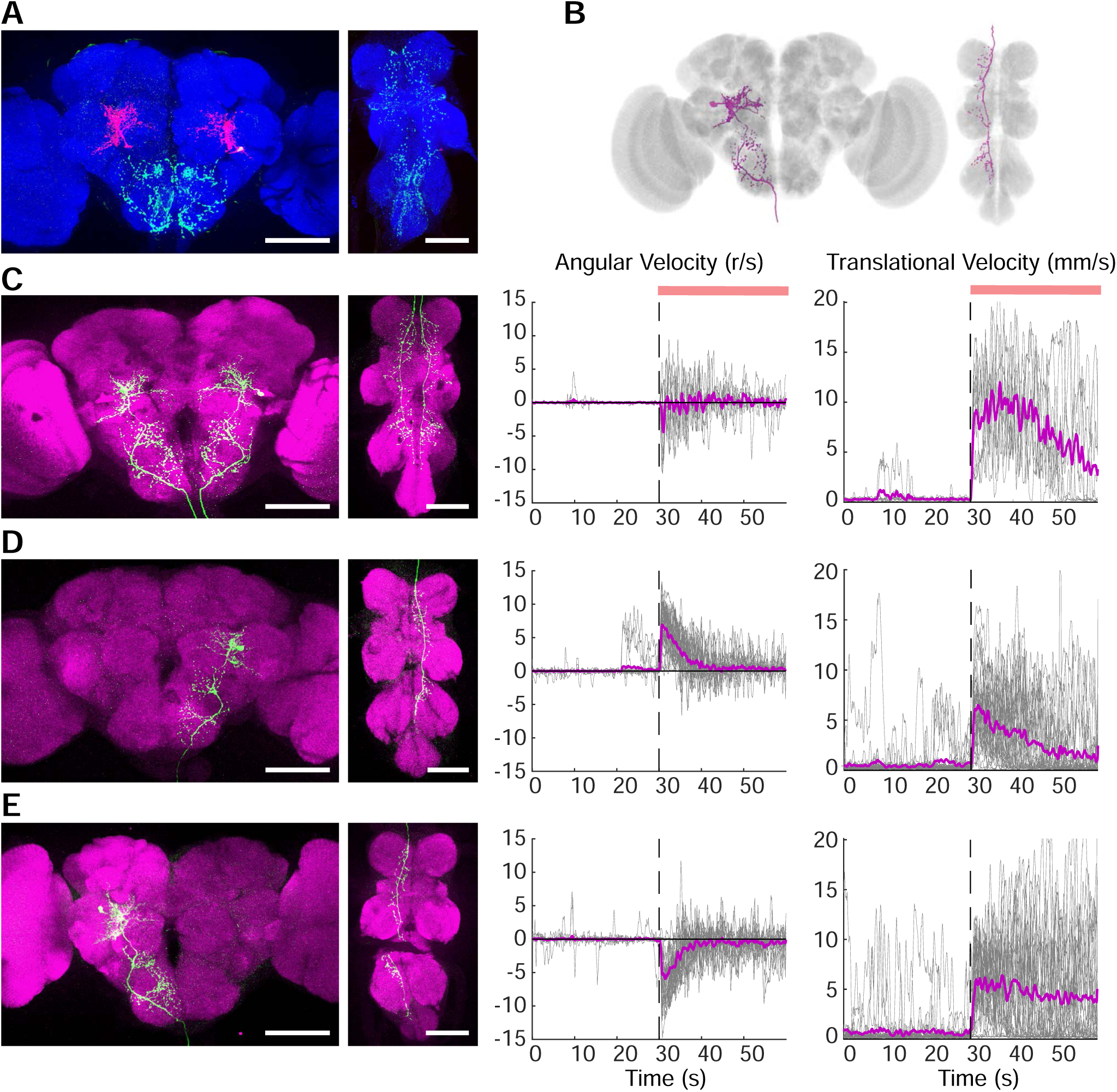
P9 activation triggers forward walking with ipsilateral turning. (A) DenMark (magenta) and synaptotagmin (green) in P9 neurites in the central brain (left) and VNC (right) of SS01540>DenMark,Syt-GFP flies. nc82 stains neuropil (blue). Scale 100 μm. (B) P9 segmented image, mapped onto a fly brain template. (C) CsChrimson-mVenus (green) and neuropil (magenta) (left), angular velocity with positive velocity for right turns, negative velocity for left turns (middle) and translational velocity (right) for mosaic animals with CsChrimson-mVenus in P9 neurons bilaterally (3 flies, 9 trials). (D) CsChrimson-mVenus (green), angular velocity and translational velocity for mosaic animals with CsChrimson-mVenus in right P9 neurons (10 flies, 30 trials). (E) CsChrimson-mVenus (green), angular velocity and translational velocity for mosaic animals with CsChrimson-mVenus in left P9 neurons (9 flies, 27 trials). Individual trials (grey), mean (magenta), light ON (red bar). See Figure S2 for characterization of an independent P9 split-Gal4 line.

### P9 neurons receive inputs from courtship promoting neurons and visual projection neurons

As P9 descending neurons provide a direct link between the central brain and the VNC and initiate ipsilateral forward turns, we hypothesized that P9 might be a key relay in a sensorimotor pathway involved in steering toward attractive objects. One vital behavior requiring a fly to frequently turn toward an attractive stimulus is male pursuit of a female during courtship behavior (Agrawal and Dickinson, 2019; Agrawal et al., 2014; Dickson, 2008; Kohatsu and Yamamoto, 2015). If P9 participates in object-directed walking during courtship, then we would predict that P9 might receive inputs from courtship promoting neurons. Consistent with this prediction, P9 dendrites arborize in close proximity to pC1, the master courtship promoting neurons in the fly (Bath et al., 2014; Hoopfer et al., 2015; Inagaki et al., 2014; Koganezawa et al., 2016; Kohatsu et al., 2011a; von Philipsborn et al., 2011) (Figures 3A and S3A). To test if P9 receives inputs from pC1, we monitored responses in P9 by GCaMP6s calcium imaging while optogenetically activating the pC1 cluster with CsChrimson (Koganezawa et al., 2016). This produced reliable calcium transients in P9 (Figures 3B and 3C), indicating that P9 neurons likely receive inputs from pC1.

**Figure 3:**
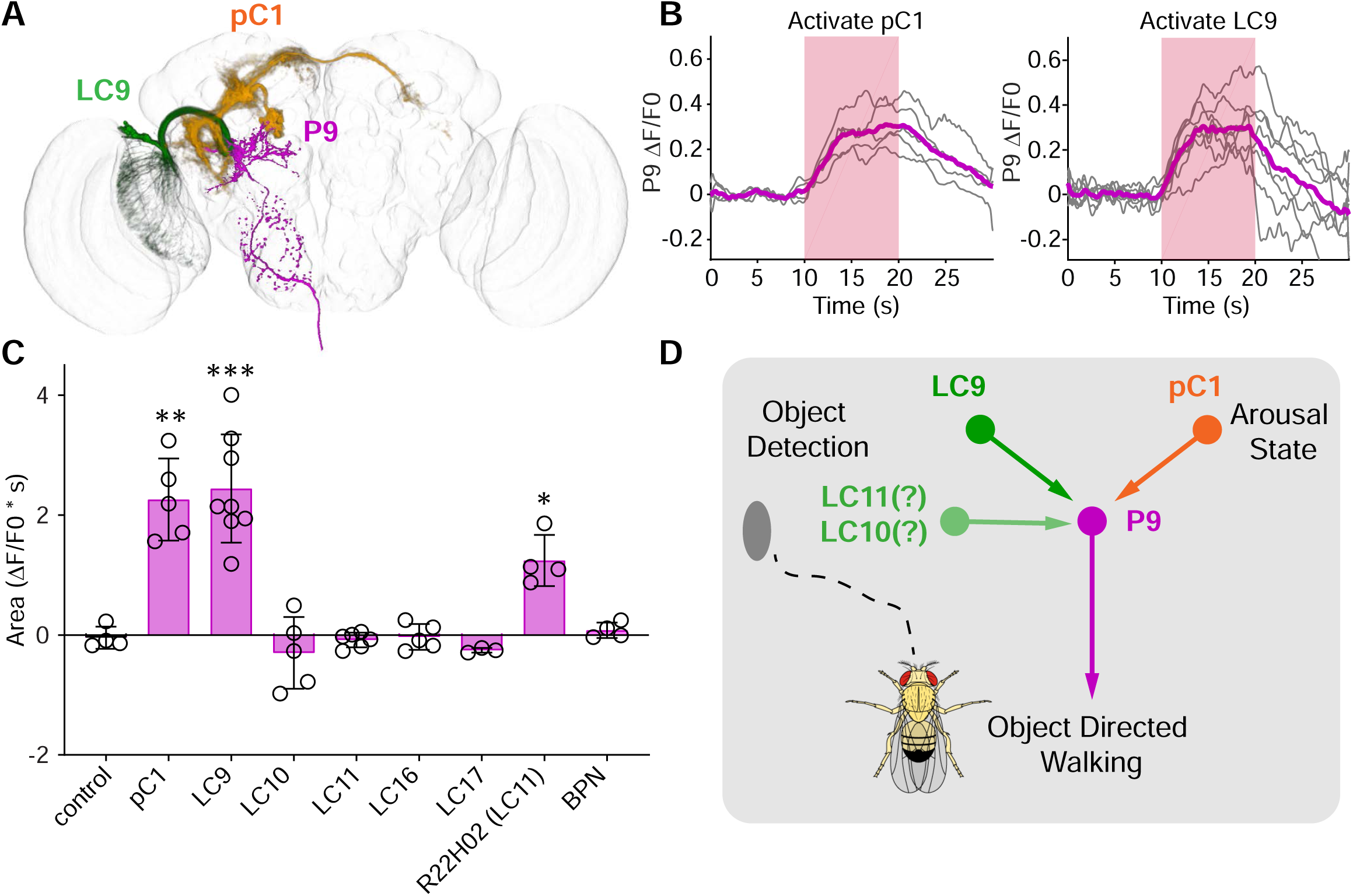
P9 neurons are activated by courtship promoting neurons and visual projection neurons. (A) P9, pC1 and LC9 segmented neurons registered onto a template brain. See Figure S3 for anatomical overlap in single brains. (B) GCaMP6s responses (ΔF/F) in P9 soma (using P9-LexA, lexAop-GCaMP6s) upon stimulation of pC1 (left) or LC9 (right), using UAS-CsChrimson-mVenus and R71G01-Gal4 for pC1 or LC9 split-Gal4. Trial averaged traces (grey), mean (magenta), optical stimulation (pink). n=5-8 flies/genotype. (C) P9 responses (area under average trial deltaF/F0 curve during light ON period) upon optical stimulation of candidate upstream neurons. n=3-8 per genotype, non-parametric Mann-Whitney test compared to no Chrimson control, *p<0.05, **p<0.01, ***p<0.001. (D) P9 receives inputs from LC9 and potentially LC10, LC11 as well as PC1, suggesting that it participates in object-directed tracking during courtship.

In addition to pC1, P9 dendrites arborize near the outputs of lobular columnar (LC) neurons (Figures 3A and S3B) that encode visual features relevant for specific behaviors. Recent studies have implicated specific LC neurons in object tracking (LC11 (Keleş and Frye, 2017)), courtship (LC10 (Kohatsu et al., 2011b; Ribeiro et al., 2018), LC9, LC16, LC17 (McKinney and Ben-Shahar, 2018; Ribeiro et al., 2018)) and figure detection (LC9 and LC10 (Aptekar et al., 2015)). To test the hypothesis that P9 receives visual inputs important for object tracking/courtship, we tested for functional connectivity between the five candidate LC neurons and P9. We used the LC split-Gal4 collection (Wu et al., 2016) for CsChrimson activation of LC candidates while monitoring P9 GCaMP6s responses. LC9 activation, using the highly specific split-Gal4 driver, produced reliable responses in P9 (Figures 3B and 3C). Moreover, immunostaining revealed that LC9 projections closely overlap with P9 dendrites (Figure S3B), consistent with direct connections. The other tested LC split-Gal4 drivers did not elicit responses in P9 (Figure 3C). Stronger activation of LC11 with a sparse LC11-Gal4, which did not show expression in other LCs, also elicited reliable responses in P9 (Figure 3C), indicating that P9 is likely to receive input from LC9 and potentially LC11.

Taken together, these functional connectivity studies demonstrate that P9 receives information from LC neurons that participate in object detection and/or tracking and pC1. This suggests that P9 might function as a node for integration of visual features and courtship cues to drive a complex motor program to pursue females during courtship (Figure 3D).

### P9 neurons are essential for object directed walking during courtship

To directly test the hypothesis that P9 participates in orienting a male toward a female during courtship, we blocked the synaptic output of P9 by selective expression of tetanus toxin (TeTx) in males and examined male courtship behavior. P9 silenced males showed overt courtship defects, with only a fraction of flies successfully copulating in a 10-minute assay (Figures 4A and 4B). Blocking transmission of potential P9 inputs, pC1 and LC9, also significantly reduced copulation (Figures 4B and S3C). To examine whether P9 silenced males show defects in following females, we evaluated the female’s position with respect to the male and the time spent in close proximity to the male pre-copulation (Figures 4C and 4D). This revealed that P9 silenced males showed significant defects in pursuing females. Silencing LC9 also decreased following, while silencing other candidate upstream neurons had no significant effect (Figure 4D).

**Figure 4:**
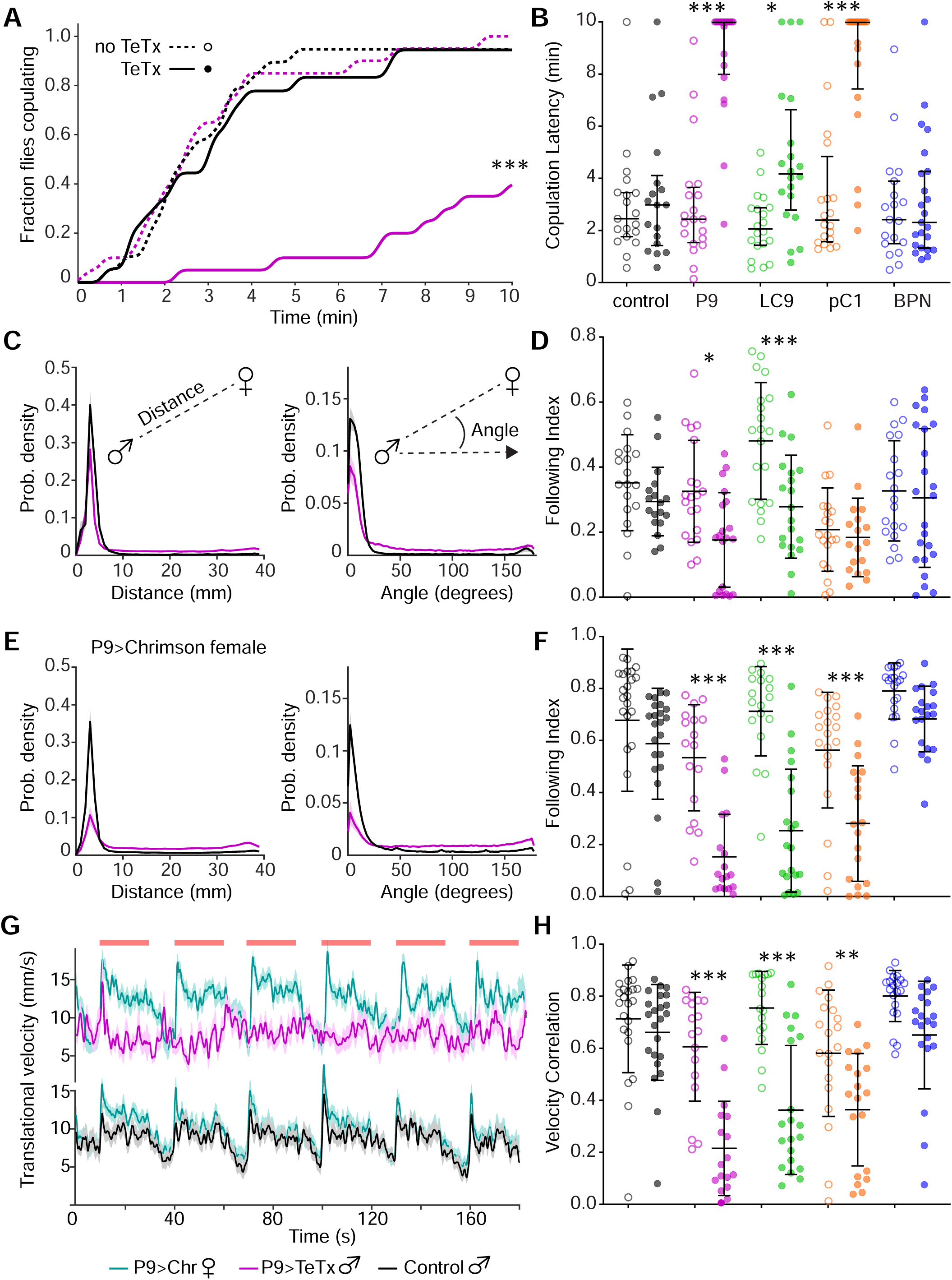
P9 is required for males to track females during courtship. (A) Fraction of males copulating with Canton S virgins over 10 minutes. Gal4 controls (dotted lines; open circles for other panels), Gal4, UAS-TeTx flies (solid lines; filled circles for other panels), n=20-28 flies/genotype, Fisher’s Exact Test, t = 10 min, ***p<0.001. Genotypes for all panels are color coded as B. (B) Copulation latency. Lines indicate median and interquartile range. n=20-28 flies/genotype, Kruskal-Wallis test and Dunn’s multiple comparison, *p<0.05, ***p<0.001. (C) Probability density showing distance (left) or angle (right) between male and female, mean ± SEM. (D) Following Index per courting pair. Lines indicate median and interquartile range. n=20-28 flies/genotype, ANOVA and Sidak’s tests, *p<0.05, ***p<0.001. (E) Probability density showing distance (left) or angle (right) between male and P9>CsChrimson “remote-controlled” female, mean ± SEM. (F) Following Index per courting male and P9>CsChrimson female. n=17-23 flies/genotype, ANOVA and Sidak’s tests, ***p<0.001. (G) Translational velocity of P9>TeTx males and P9>CsChrimson females, or control males and P9>CsChrimson females, showing mean (dark) and SEM (shading), n = 17-23 pairs. (H) Correlation between male and female velocities per courting pair, for males courting P9>CsChrimson females. n = 17-23 pairs, t-test with Sidak’s multiple comparison corrections, **p<0.01, ***p<0.0)1. See Figure S3 for additional characterization of P9 courtship phenotypes. See Figure S4 for gait analysis of P9>TeTx flies.

Most control flies copulated soon after the assay started and showed short mate pursuit sequences, making it challenging to evaluate the tracking ability. To encourage the male to perform long and complex tracking behavior, we replaced the target female with a virgin female expressing CsChrimson in P9 neurons. The “remote-controlled” female was intermittently triggered to perform sudden forward walking and turning via P9 activation throughout the assay, making it a challenging target. Control males engaged in long following sequences chasing remote-controlled females (Figures 4E, 4F and Video S4). Moreover, control males exhibited rapid velocity changes that were synchronized with the artificially evoked sudden velocity changes of the remote-controlled females (Figures 4G, 4H and Video S4). P9 silenced males showed significantly reduced following of the remote-controlled females (Figures 4E and 4F). In addition, the velocities of P9 silenced males showed little correlation to the velocities of the remote-controlled females (Figures 4G and 4H), although the males walked at similar average velocities as controls (Figure S3D). Blocking activity of neurons functionally upstream of P9, including LC9 and pC1, also resulted in courtship following deficits (Figures 4F, 4H and S3E). To test male pursuit towards non-manipulated females, we designed flat, low ceiling arenas, which enhanced the repertoire of female locomotion. Results in this assay were similar to the remote-controlled female assay (Figures S3F-H), demonstrating that P9 and upstream neurons are required for following a female during courtship.

Although the average walking velocities of P9 silenced flies were similar to controls, we wondered if defects in stepping or inter-leg coordination were responsible for the poor object pursuit of these flies. To test this, we analyzed the walking pattern of P9 silenced flies in fine detail via high speed recordings and deep learning based automated leg position tracking (Mathis et al., 2018). This analysis showed that P9 silenced flies have normal stepping parameters and inter-leg coordination (Figure S4). Thus, the behavioral deficits observed in courtship do not reflect uncoordinated walking.

Taken together, these studies demonstrate a specific role for P9 DNs in visually guided pursuit behavior during courtship. Neurons functionally upstream of P9, including LC9 and pC1, also showed courtship following deficits, supporting the notion that they contribute to an object tracking pathway that is likely gated by courtship state. While P9 inactivation precludes a male’s ability to follow a female, other aspects of locomotion are unaffected. Thus, the studies of P9 reveal a specific locomotor program for object-directed walking.

### Bolt Protocerebral Neurons (BPNs) initiate straight forward walking

Because P9 impacts a specific walking program, this suggests that there may be independent pathways for different forward walking behaviors, for example, object-directed steering versus random search behavior. To examine if there are multiple independent pathways that drive different walking behaviors, we returned to the results of our walking initiation screen and characterized the walking behavior of the second candidate walking initiation line, SS01587 (Namiki et al., 2018), which induced straight forward walking without turns upon activation (Figures 1E, 1F, S1D and S1E).

SS01587 labels DN0p28 and 7 other neuronal types (Figure S5A). To determine the neurons in SS01587 causal for walking initiation, we used mosaic strategies to restrict the number of neurons expressing CsChrimson in SS01587 and screened for mosaic animals that initiated walking in grooming flies upon transient activation. Comparing CsChrimson expression in the brains of walking and non-walking individuals revealed that activation of a single neuronal type in SS01587 is correlated with walking initiation (Figure S5B). This neuronal type, which we name Bolt Protocerebral Neurons (BPNs), is a cluster of ~10 cells per hemisphere, with cell bodies located near the posterior superior margin of the fly brain (Figure S5C). To obtain specific, reproducible genetic access to BPNs, we generated two new split-Gal4 reagents that label BPNs (Figures 5A and S5D). Both lines triggered robust walking initiation in grooming flies upon optogenetic activation (Figures 5B and S5E-H). As with SS01587, these flies initiated straight forward walking (Figure S5F).

**Figure 5:**
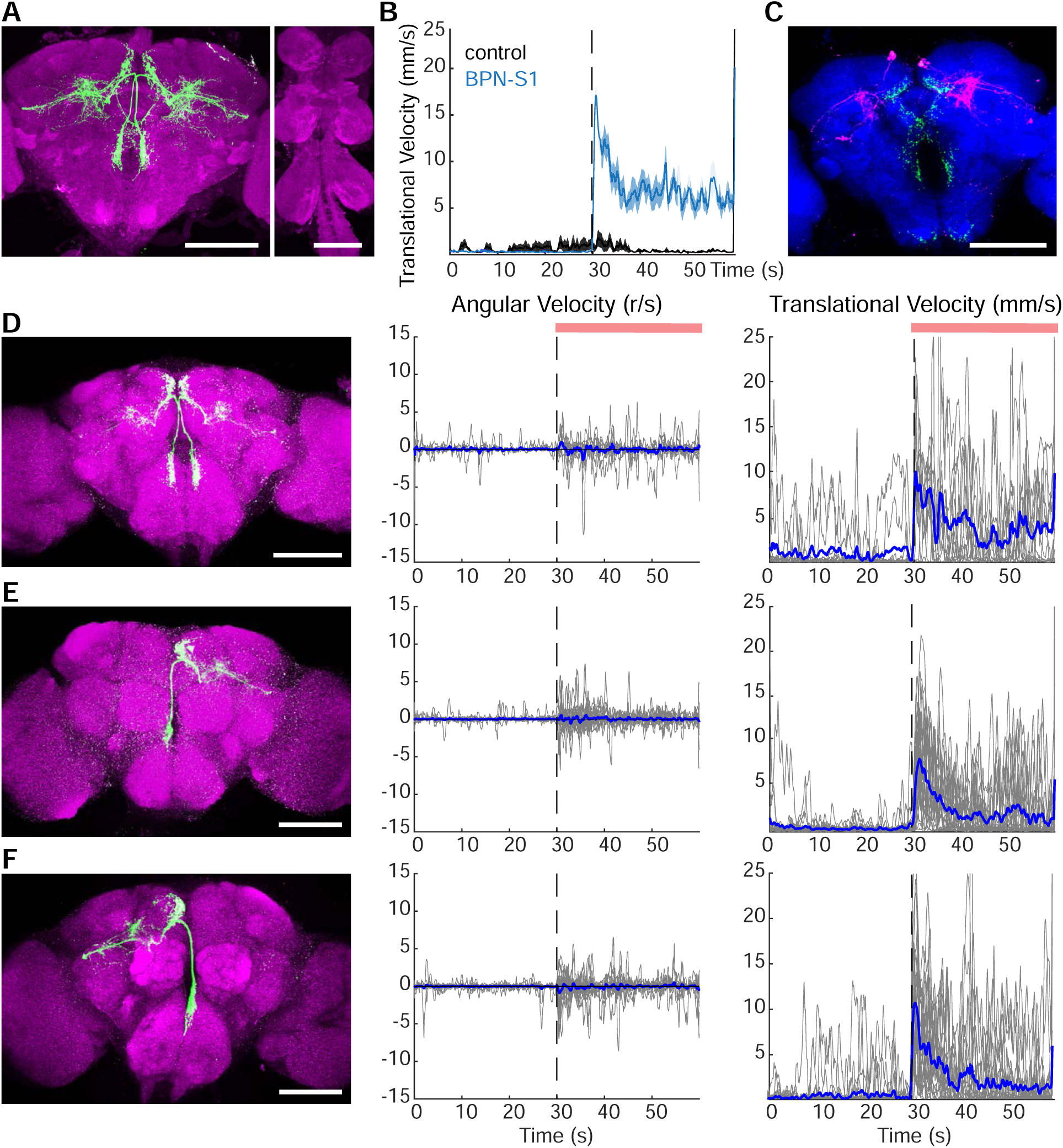
BPNs trigger forward walking with no turning bias. (A) BPN-S1 split-Gal4 line, showing specific BPN expression (green), nc82 stains neuropil (magenta). (B) Translational velocity upon BPNS1 split-Gal4 activation with CsChrimson in grooming flies. n=8 flies, 24 trials, mean ± SEM. (C) DenMark (magenta) and synaptotagmin (green) labeling in BPN. nc82 stains neuropil (blue). Scale 100 μm. (D) CsChrimson-mVenus (green) (left), angular velocity plot with positive velocity for right turns, negative velocity for left turns (middle) and translational velocity (right) for mosaic animals with CsChrimson-mVenus in BPNs bilaterally (4 flies, 12 trials). (E) CsChrimson-mVenus (left), angular velocity (middle) and translational velocity (right) for mosaic animals with CsChrimson-mVenus in right BPNs (8 flies, 24 trials). (F) CsChrimson-mVenus (left), angular velocity (middle) and translational velocity (right) for mosaic animals with CsChrimson-mVenus in left BPNs (8 flies, 24 trials). Graphs in D-F show individual activation trials (grey), mean (magenta), light ON (red bar). See Figure S5 for additional characterization of 2 BPN split-Gal4 lines.

BPN dendrites arborize extensively in the lateral protocerebrum and axons project along the contralateral midline towards the subesophageal zone (SEZ) (Figure 5C). If BPNs receive inputs regarding spatially localized sensory stimuli, then one prediction would be that unilateral activation of BPNs would produce a turning component to the induced walking, similar to P9 unilateral activation. However, unlike P9, unilateral activation of BPNs did not confer a turning bias and instead evoked straight forward walks. Comparing translational velocity and angular velocity for animals containing CsChrimson in the left BPNs, right BPNs, or both revealed no differences (Figures 5D-F). Thus, BPN activation produces straight, forward walking without a turning bias.

The activation phenotypes of BPNs and P9 are distinct, with P9 activation eliciting walking with ipsilateral turns, and BPNs eliciting straight, forward walks, suggesting that they are components of two independent walking initiation pathways. Consistent with this, activating SS01587 (BPN) and SS01540 (P9) simultaneously increased walking velocities and distance travelled compared to activation of each line alone (Figure S5I). In addition, silencing BPN did not affect copulation or mate pursuit in all courtship assays (Figure 4 and S3), arguing that BPN is not essential for object directed walking. Finally, as BPN is a higher brain neuron and P9 is a descending neuron, we tested whether BPN is upstream of P9, by CsChrimson mediated BPN activation and P9 GCaMP6s imaging, and found that BPN did not activate P9 (Figure 3C). Taken together, these studies demonstrate that BPN and P9 participate in independent walking initiation pathways, consistent with their different behavioral phenotypes.

### BPN activity correlates with high velocity straight, forward walking

We examined BPN activity during spontaneous walking behavior to evaluate its response properties *in vivo*. BPN activity was monitored by GCaMP6s imaging of BPN soma while flies were walking on an air-supported ball and their locomotion was recorded (Figure 6A). Interestingly, mean BPN activity was strongly correlated to walking velocity only in a subset of walking bouts and trials. Not all BPN soma showed measurable activity; these were not included in analyses.

**Figure 6:**
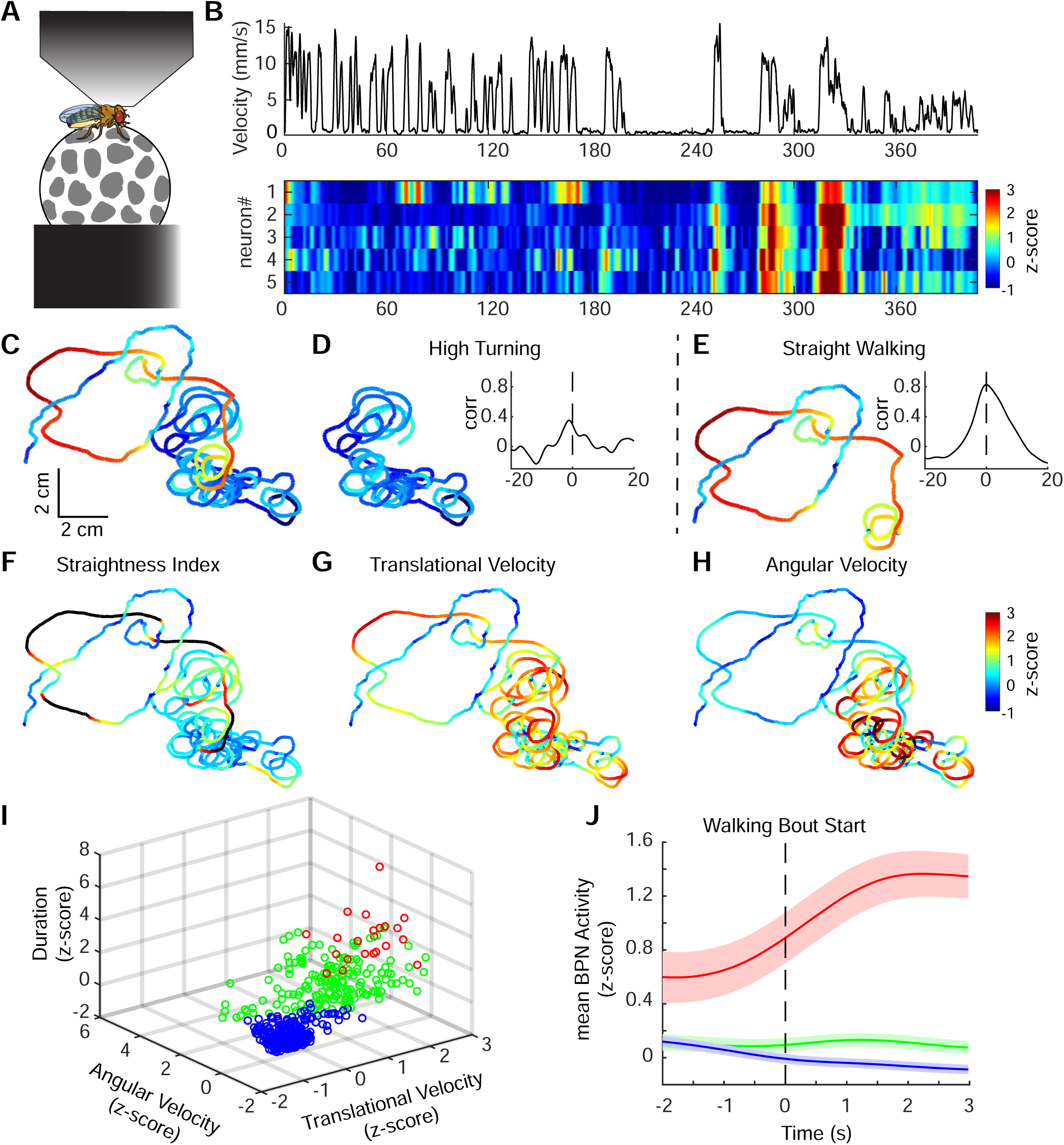
BPNs activity correlates with long, straight forward walks. (A) in-vivo imaging schematic of a fly walking on a ball. (B) top: translational velocity of one fly during an imaging session, bottom: time-locked imaging readout of 5 BPN soma in the same fly, shown as z-scored GCaMP fluorescence. (C-E) Trajectory of the fly in B color coded for mean BPN activity (z-score color-scale as B for the entire session (C), or first (D) or second half (E). Velocity versus mean BPN activity correlation plots for the first (D graph) and second half (E graph) of the imaging session. (F-H) Trajectory of fly in B color coded with straightness index (F), translational velocity (G) and angular velocity (H). (I) 3D subspace (velocity, angular velocity and walking duration.) representation of unbiased k-means clustering of 589 walking bouts across 3 flies and 5 imaging sessions. (J) BPN activity (mean ± SEM) for each cluster aligned to walking bout start (dotted line). Cluster colors as in I. See Figure S6 for additional analysis of BPN activity.

To examine when BPNs are recruited by the fly, we mapped BPN calcium activity onto the walking trajectory. In one illustrative example, the walking trajectory was initially comprised of tight turning bouts and low BPN activity uncorrelated to translational velocity (Figures 6B-D). Subsequently, the trajectory was comprised of longer straight walking bouts and high BPN activity correlated to translational velocity (Figure 6E), indicating that BPNs might be specifically recruited during straight walking. Overall, BPN activity was high when translational velocity (V_T_) was high and angular velocity (V_A_) was low (Figures 6F-H and S6A). Additionally, BPN activity was correlated to duration of walking bouts (T_W_) (Figure S6A). A linear regression model using V_T_, V_A_ and T_W_ as predictors of BPN activity showed positive regression coefficients for V_T_ and T_W_, but negative for V_A_ (Figure S6B) corroborating that BPNs are preferentially active during straight walking bouts.

To obtain an unbiased view of how BPN activity correlates with walking, data across 3 flies and 5 imaging sessions, amounting to 589 walking bouts, was pooled for unbiased k-means clustering in a 4D space of [mean BPN activity, V_T_, V_A_, T_W_]. One cluster corresponds to high BPN activity, V_T_ and T_W_, but low V_A_ (red cluster, Figures 6I and S6D-F). This cluster is comprised of high velocity, long duration, low turning bouts and represents the majority of instances when BPNs were highly active during walking initiation (Figures 6J and S6D-F). In contrast, the other two clusters are comprised of low BPN activity and high angular velocity bouts (Figures 6I, 6J and S6D-F). Thus, this independent analysis confirms that BPN activity specifically increases during fast, long, straight walks.

### BPN activity reciprocally regulates straight, forward walking

If BPN activity drives straight, forward walking without turns, then one prediction would be that artificially increasing BPN activity would increase walking bout duration and translational velocity but not angular velocity. To test this, we expressed CsChrimson in BPNs, stimulated BPNs with a range of light stimulation frequencies (from 10-150 Hz) and monitored walking behavior in flies that were grooming. As predicted, the total distance, walking duration and translational velocity of walking increased with stimulation frequency, scaling linearly as a function of activation frequency (Figure 7A). Moreover, in line with our observation that BPN activity did not increase during high turning states, optogenetically increasing BPN activity did not produce a similar increase in angular velocity.

**Figure 7:**
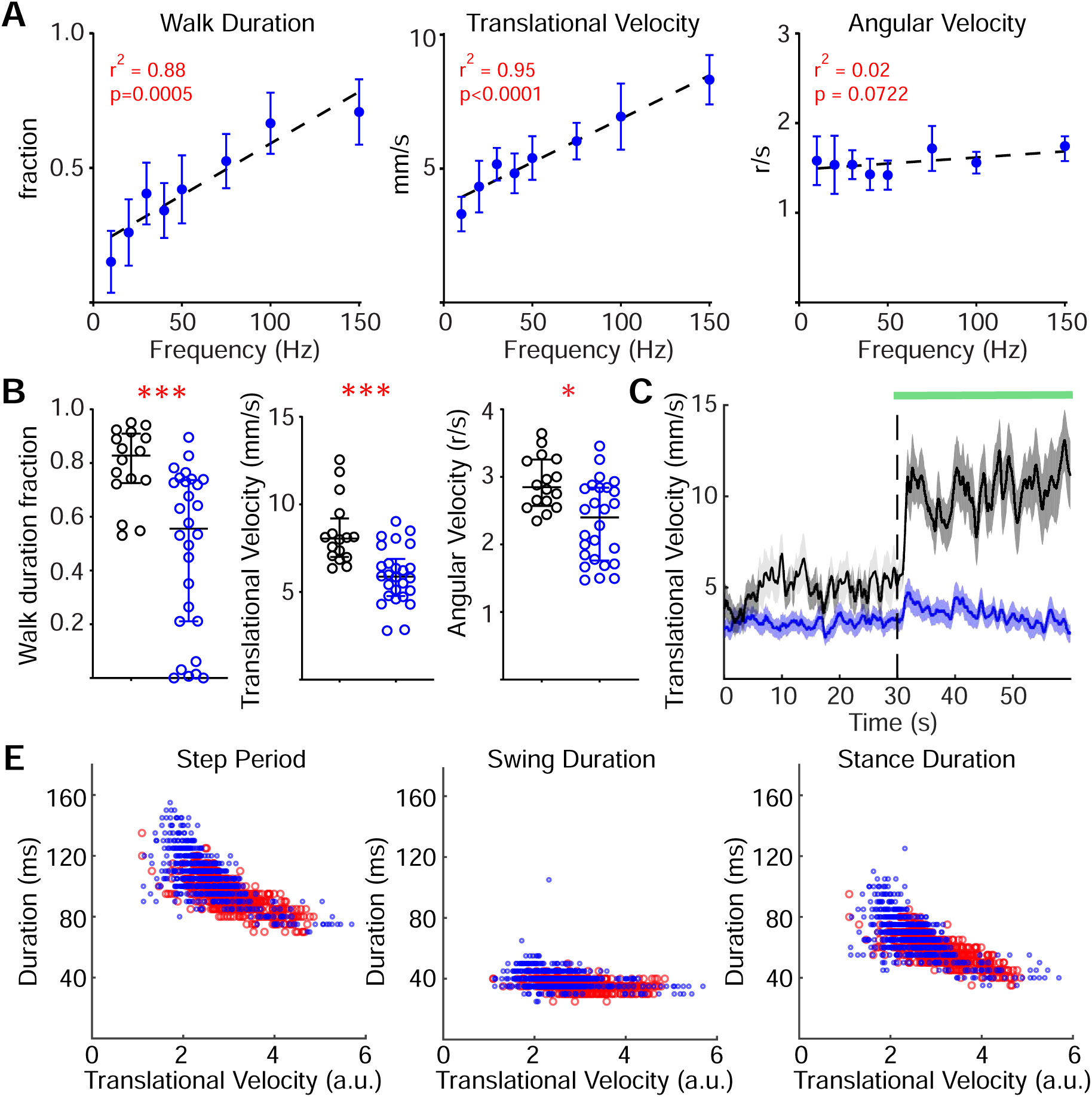
BPN activity levels reciprocally regulate straight, forward walking. (A) Walk duration (left), translational velocity (middle) and angular velocity (right) as a function of optogenetic activation (LED frequency) for BPN>CsChrimson flies. n=20 flies, mean ± 95%CI, linear regression line (dotted line) for r^2^ values shown. (B) Walk duration (left), translational velocity (middle) and angular velocity (right) of controls (black) and BPN>TeTx flies (blue). n= 6-28, Mann-Whitney, *p<0.05, ***p<0.001. (C) Trial averaged translational velocity during green light stimulation for controls and BPN>TeTx flies. n=16-28, mean ± SEM, green bar indicates light on. (D) Middle leg step period (left), swing duration (middle) and stance duration (right) as a function of translational velocity of controls (red) and BPN>TeTx (blue), n=283-314 steps from 4-5 flies, t test. See Figure S7 for additional high-resolution quantification of BPN silenced flies.

Conversely, to test whether BPNs are necessary for fast, straight walks, we silenced BPN outputs by expressing TeTx. Freely walking flies with synaptic transmission blocked in BPNs showed a small but significant reduction in walking duration, translational velocity and angular velocity (Figure 7B). To test if flies might require BPN activity for a sudden increase in walking speed, flies initially kept in the dark were suddenly exposed to bright green light. The light stimulated high speed walking in control flies, whereas BPN silenced flies showed a smaller velocity change and lower maximum velocities (Figure 7C). Critically, high resolution analyses of walking gait in freely walking flies revealed that BPN-TeTx flies were not uncoordinated (Figure S7) but instead walked slowly as a result of an increased step period, specifically due to an increased stance duration (Figure 7E). Previous work has shown that flies typically vary walking speed by varying stance duration (DeAngelis et al., 2019; Wosnitza et al., 2013). Therefore, the gait of BPN silenced flies resembles that of slow walking flies. In contrast, P9 silenced flies did not show decreased translational velocity or increased stance duration (Figure S4). Taken together, these experiments demonstrate an essential role for BPNs in driving walking at high translational velocity and further demonstrate that P9 and BPN drive distinct motor programs.

## Discussion

This study identifies two classes of central neurons sufficient to initiate coordinated forward walking in the fly, representing a significant step forward in studies of locomotor control. P9 DNs receive inputs from courtship promoting neurons and visual projection neurons and drive forward walking with an ipsilateral turning component. Moreover, P9 is required during object directed walking in the context of courtship, whereas BPN is not required for object directed walking in this context. Instead, BPN activity is correlated with straight, fast walking bouts and manipulating activity of these neurons modifies walking speed and duration without affecting turning. Thus, the characterization of these neurons reveals two distinct pathways that initiate two different forward walking programs in different behavioral contexts.

### Distinct walking initiation commands initiate distinct walking programs

P9 and BPN uncover separate brain pathways for object-directed forward steering and fast straight forward walking, respectively. The existence of these independent walking initiation pathways argues that specific brain inputs can drive distinct and complete walking programs in a context specific manner. This may be achieved by driving a common downstream walking circuit that is modulated to generate specific walking modes, as suggested by neuromechanical models for directional control of walking (Guschlbauer et al., 2012; Tóth and Daun, 2019; Tóth et al., 2012). Alternatively, these pathways may recruit distinct downstream premotor circuits, each generating a distinct motor pattern. Examining the downstream circuits of these walking initiation pathways will help unravel the mechanisms of these distinct walking initiations.

Complex motor patterns observed in natural behaviors likely result from combined activity of populations of descending commands from the brain (Pearson, 1993). Interactions among DNs may bring about a concerted change in the functional state of nerve cord circuits, as seen in crawling circuits in *C. elegans* (Piggott et al., 2011) and *Drosophila* larvae (Carreira-Rosario et al., 2018). Therefore, in natural behaviors, P9 and BPN pathways may be active in parallel with other descending signals. An interesting question is whether a population of descending commands generates a motor output more complex than the sum of its constituents. Genetic access to characterized walking initiation neurons (P9 and BPN) will now enable examination of this question and will guide future experiments aimed at elucidating how brain neurons influence walking control.

### The P9 walking program provides steering control during object tracking

From an activation screen of approximately one third of all DNs in the fly (Namiki et al., 2018), the P9 DNs were the only DNs to initiate walking. These results argue that very few single DNs encode forward walking initiation. Similarly, a previous unbiased behavioral screen revealed MDN as the only DN sufficient to drive backward walking (Bidaye et al., 2014). While DN combinations likely elicit and modulate walking, these studies argue that P9 neurons have a privileged role as a DN class triggering a complete forward walking program.

Previous studies have shown the strong importance of vision in pursuing a potential mate during courtship (Cook, 1980; Kohatsu and Yamamoto, 2015; Markow and Manning, 1980; Ribeiro et al., 2018). Male flies initiate walking towards any object with visual characteristics that grossly match that of a potential mate (Agrawal and Dickinson, 2019), indicating that visual information may directly influence walking control neurons. Here, we showed that P9 receives inputs from LCs that serve as feature detectors, arguing that P9 participates in a shallow pathway from visual detection to motor command. Direct connections between visual projection neurons and DNs are also seen in the Giant Fiber escape pathway (Klapoetke et al., 2017; von Reyn et al., 2017; Strausfeld and Bassemir, 1983) and in visually guided flight control (Strausfeld and Gronenberg, 1990; Suver et al., 2016), suggesting a common circuit motif to minimize response times when rapid action is required.

In addition to visual inputs, P9 is also activated by pC1 neurons, master regulators of courtship behavior that integrate olfactory, pheromonal, and auditory cues and drive a complex courtship sequence (Clowney et al., 2015; Coen et al., 2016; Kallman et al., 2015; Kohatsu and Yamamoto, 2015; von Philipsborn et al., 2011). pC1 activated flies show enhanced object pursuit behavior that persists for several minutes (Agrawal et al., 2014; Bath et al., 2014; Inagaki et al., 2014; Kohatsu and Yamamoto, 2015; Rezával et al., 2016), suggesting strong potentiation of the downstream locomotor control circuit. Our findings reveal a circuit configuration where pC1 may gate/potentiate the LC-P9 sensorimotor loop (Figure 3E), leading to context-specific object directed walking. Consistent with this, recent studies demonstrated that LC10 neurons participate in object tracking during courtship and proposed that LC10 and pC1 converge on downstream targets to drive object directed walking. Although LC10 did not activate P9 (Figure 3D), it is possible that its effect on P9 is gated by pC1 activity. Similar to the P9 pathway, state-dependent gating of sensorimotor loops in *Drosophila* was recently demonstrated for a visually evoked landing circuit (Ache et al., 2019), suggesting a general circuit architecture for context-dependent, modular regulation of sensory-driven responses.

Although we defined a role for P9 in object pursuit during courtship behavior, it is unlikely that P9 neurons are the only DNs involved in walking control during courtship, as males with P9 transmission blocked are able to track females for short periods. In addition, we have no evidence that P9 is active only during courtship behavior and have not explored the role of P9 in other steering behaviors. Indeed, cricket DNs implicated in walking control have been shown to be responsive to multiple sensory stimuli in a state-dependent fashion (Staudacher, 2001).

### Bolt Protocerebral Neurons (BPNs) drive fast, straight forward walking

The ability of BPNs to drive high speed, straight forward walks (or “sprints”) inspired the name “Bolt” neurons, given Usain Bolt’s unmatched sprinting records. BPNs display an anatomy and function different from other neurons implicated in insect walking control. Their widespread dendritic fibers in the higher brain suggest that they might integrate multiple sensory inputs and may represent higher order regulators of walking, akin to pC1 neurons for courtship control. Unilateral BPN activation induced straight forward walking, in contrast to turning phenotypes elicited by unilateral activation of other walking initiation neurons (P9, MDN and cricket DNs) (Böhm and Schildberger, 1992; Sen et al., 2017; Zorović and Hedwig, 2013). The BPN phenotype suggests that a neural pathway operating via unidentified DNs elicits straight forward walking. BPNs also likely function in speed control, as increasing activity increased walking speed.

The BPN pathway shows remarkable similarity to a recently described mouse MLR-caudal brainstem circuit that promotes high-speed walking: in both cases, unilateral activation induces walking initiation without a turning bias, activation strength correlates with walking speed, and reduced activity causes specific defects in high speed walking (Capelli et al., 2017, Caggiano et al, 2018). These similarities suggest that high-speed walking is executed by specialized walking circuits that serve an essential function across different organisms.

Although BPNs are necessary for fast walking, the ability of BPNs to drive different walking speeds at different stimulation frequencies, provides an opportunity to examine downstream mechanisms for speed control. Recent studies in zebrafish (Ampatzis et al., 2014; Song et al., 2018) show a gradient of recruitment of distinct premotor circuits at increasing swimming speeds. In *Drosophila*, recent work (Azevedo et al. 2019) has characterized distinct motor neurons recruited in a similar manner as leg movements accelerate. Examining how BPNs recruit these motor programs in an intensity dependent manner will help illuminate the mechanism of walking speed control in *Drosophila*.

### BPNs and P9 define novel pathways for walking control

The mammalian walking control circuit has been explored at several hierarchical levels, including the motor cortex, hypothalamus, basal ganglia, cerebellum, brainstem and spinal cord (Arber and Costa, 2018). Extensive studies have provided a framework for how different types of locomotion are executed and suggest the recruitment of different brainstem subcircuits (Klaus et al., 2017; Li et al., 2018).

In contrast to mammalian locomotor studies, invertebrate walking control has been mainly investigated at the level of the nerve cord circuits and DNs (Bässler and Büschges, 1998; Bidaye et al., 2018; Tuthill and Azim, 2018). The only higher brain structure analyzed in the context of locomotion is the central complex (CC) which has been implicated as the site for generation of an internal heading signal (Green et al., 2017, 2019; Kim et al., 2017; Seelig and Jayaraman, 2015; Turner-Evans et al., 2017). These heading signals are thought to directly influence CC output neurons involved in turning and speed control (Heinze and Pfeiffer, 2018; Martin et al., 2015; Strauss, 2002). It is unclear how CC or other higher brain areas control downstream locomotor circuits.

Our characterization of P9 and BPN pathways shows how specific brain neurons can drive distinct walking programs that are elicited in different contexts. Moreover, BPNs are higher brain neurons located outside the CC and are not involved in turning behaviors. BPNs may therefore constitute a CC independent higher brain pathway involved in non-directed high-speed forward walking and provide an important landmark in elucidating insect higher brain regions involved in walking control. Genetic access to P9 will similarly aid our understanding of a functionally distinct walking program involved in context specific object-directed walking. This study has therefore characterized walking initiation pathways at different hierarchical levels and provides an important advance in our understanding of how brain pathways switch on downstream walking control circuits.

## Acknowledgments

We thank M. Dübbert for designing the custom LED panel for transient activation experiments and ball holder for in-vivo imaging experiments. We thank J.Ho and V. Godesberg for assisting in manual annotation of behavior videos. We thank G. Card and S. Namiki for access to DN split-Gal4 collection prior to publication. We thank H. Aaron for help with designing functional connectivity imaging setup. We thank members of Scott lab, Charles Zuker and Mala Murthy for helpful feedback on the manuscript. This work was supported by grants from DFG (Bu857 to A.B.) NIH (UF1 NS107574 K.S.) and an HHMI Early Career Scientist award (K.S.).

## Author Contributions

SSB conceived the project, designed the experiments and analyzed the data under guidance from KS. SSB and AKC performed behavioral screen, expression analysis and functional connectivity experiments. ML performed courtship experiments. SSB, TB and AB performed high resolution walking analysis. SSB and YL performed stochastic activation experiments. SSB performed the in-vivo imaging experiments. SSB and KS wrote the manuscript with feedback from other authors.

## Supplementary Figure Legends

**Figure S1 (related to Figure 1):**
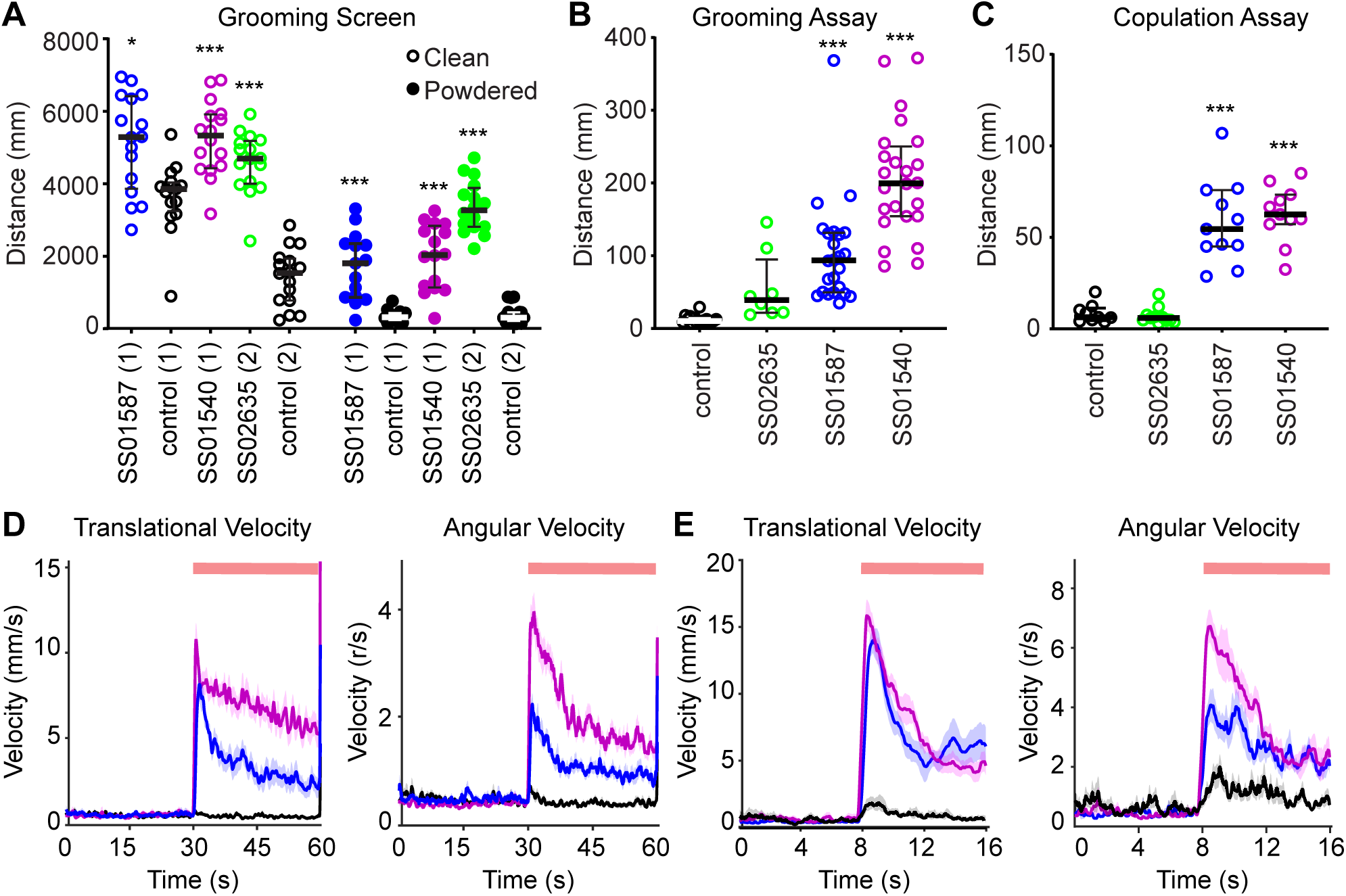
(A) Total distance traveled by clean (open circles) or powdered flies (filled circles) during the 7-minute continuous optogenetic stimulation assay. n = 15-16 per genotype, lines indicate median and interquartile range. Mann Whitney test to same day controls, *p < 0.05, ***p<0.001. (B) Distance traveled by grooming flies in Figure 1C during the light ON period (mean ± SEM), n=16-24 flies/genotype, ANOVA, Tukey post-hoc, ***p<0.001. (C) Distance traveled by copulating flies in Figure 1D during the light ON period (mean ± SEM), n=9-11 flies/genotype, ANOVA, Tukey post-hoc, ***p<0.001. (D) Translational velocity and angular velocity (mean ± SEM) for SS01540 (magenta), SS01587 (blue), and control (black) in the grooming assay. n=16-24 flies/genotype. (E) Translational velocity and angular velocity (mean ± SEM) for SS01540 (magenta), SS01587 (blue), and control (black) in the copulation assay. n=16-24 flies/genotype.

**Figure S2 (related to Figure 2).**
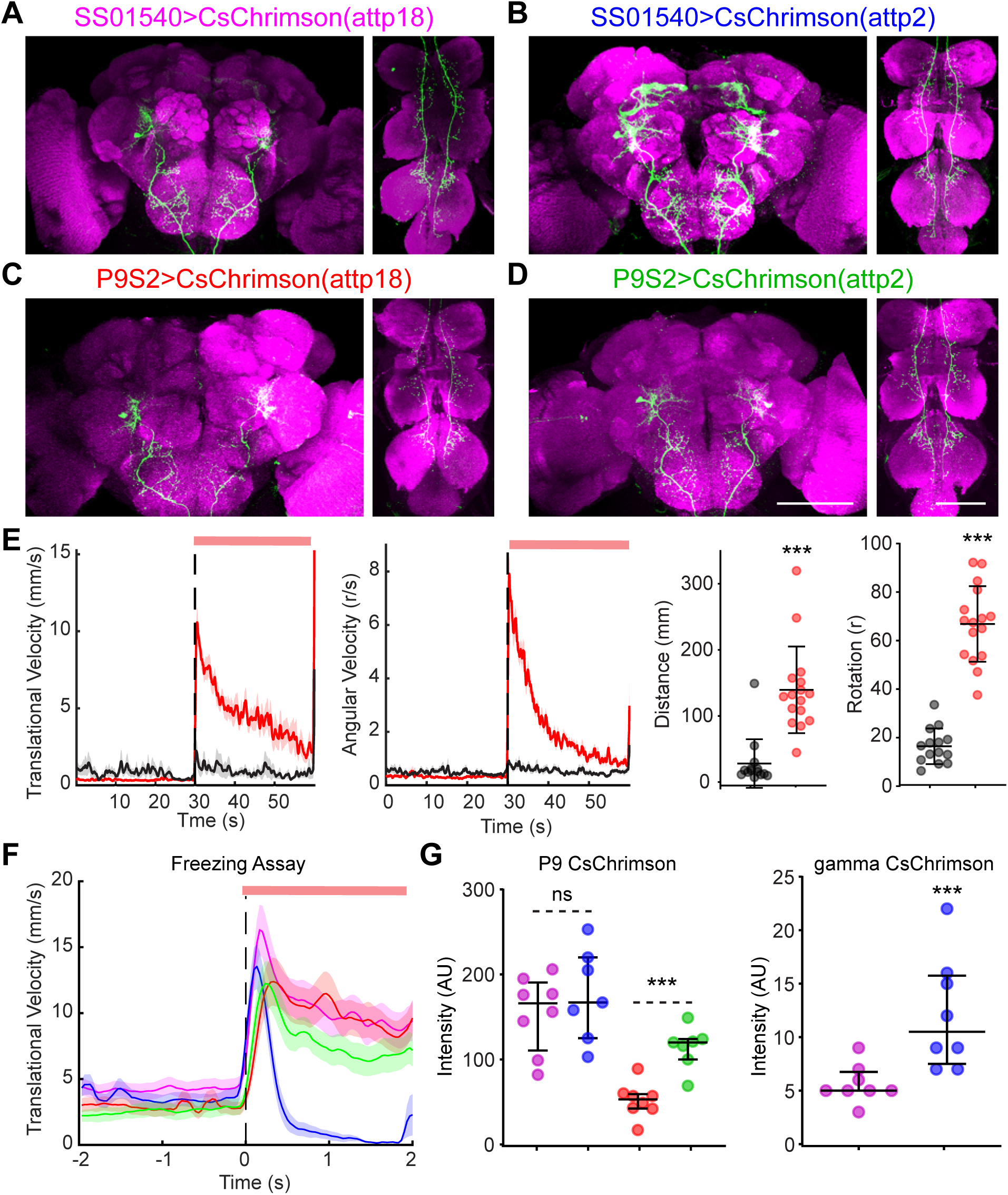
(A-D) CsChrimson-mVenus (green) and nc82 (magenta) immunostaining of age-matched female flies corresponding to different driver-reporter combinations targeting P9 neurons.Scale 100 μm. (E) Average trial translational velocity, angular velocity, distance traveled during the light ON period and rotation during the light ON period in the grooming assay for P9S2>CsChrimson(attp18) flies (red) and controls (black). n=14-16, mean ± SEM, two tailed t-test, ***p<0.001. (F) Average trial translational velocity (mean ± SEM) in the freezing assay protocol for different genotypes color coded as in A-D. (G) Quantification of CsChrimson-mVenus in age-matched explant brains in P9 neurons (left) or gamma lobe (right), color coded as in A-D. n=7-8 per genotype, non-parametric Mann-Whitney test, ***p<0.001.

**Figure S3 (related to Figure 4).**
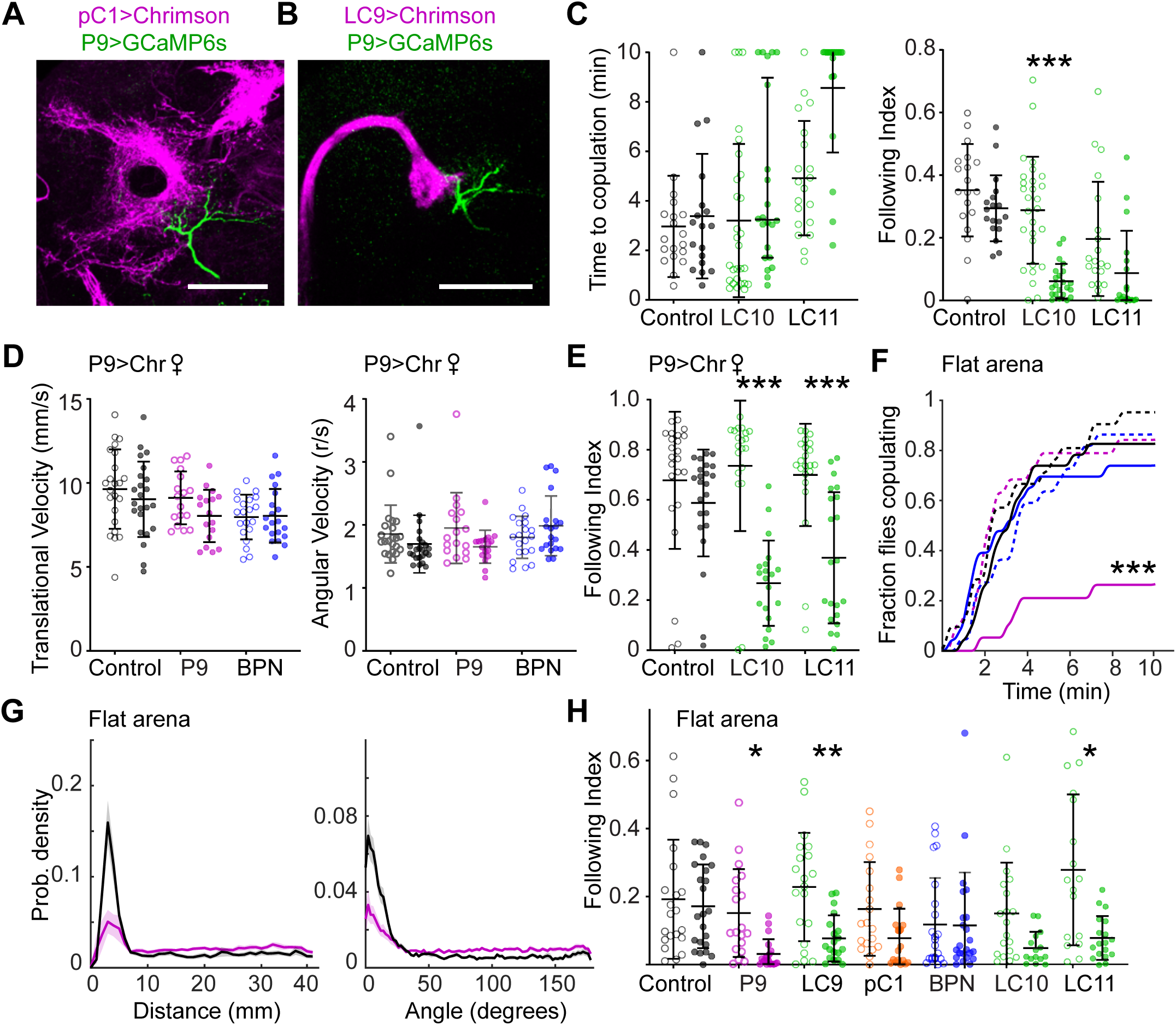
(A) pC1 lateral protocerebral ring projections (magenta) and P9 dendrites (green) in R71G01>Chrimson-tdtomato, P9>GCaMP6s flies. Scale 50 μm. (B) LC9 projections (magenta) and P9 dendrites (green) in LC9 split>Chrimson-tdtomato, P9>GCaMP6s flies. (C) Time to copulation (left) and Following Index (right) for Gal4 controls (open circles), Gal4, UAS-TeTx flies (filled circles) for all panels, n=20-28 flies/genotype, Fisher’s Exact Test (left), t = 10 min, ANOVA and Sidak’s tests (right), ***p<0.001. (D) Translational velocity and angular velocity for males courting P9>CsChrimson females. n=17-23 flies/genotype, ANOVA and Sidak’s tests. (E) Following Index (right) for Gal4 controls and Gal4, UAS-TeTx males paired with P9>Chrimson females.n=17-23 flies/genotype, ANOVA and Sidak’s tests ***p<0.001. (F) Fraction of male flies copulating with Canton S virgin females in flat arenas over a 10-minute assay duration. Dotted lines represent Gal4 controls and solid lines represent Gal4, *UAS-TeTx* flies in which synaptic transmission is blocked in specific neural classes, n=15-23 flies per genotype. ***p<0.001 Fisher’s Exact Test for time point t = 10 min. (G) Probability density showing distance (left) or angle (right) between male and female in flat arena, mean ± SEM. (H) Following Index per courting male and female in flat arena. n=17-23 flies/genotype, ANOVA and Sidak’s tests,*p<0.05, **p<0.01.

**Figure S4 (related to Figure 4).**
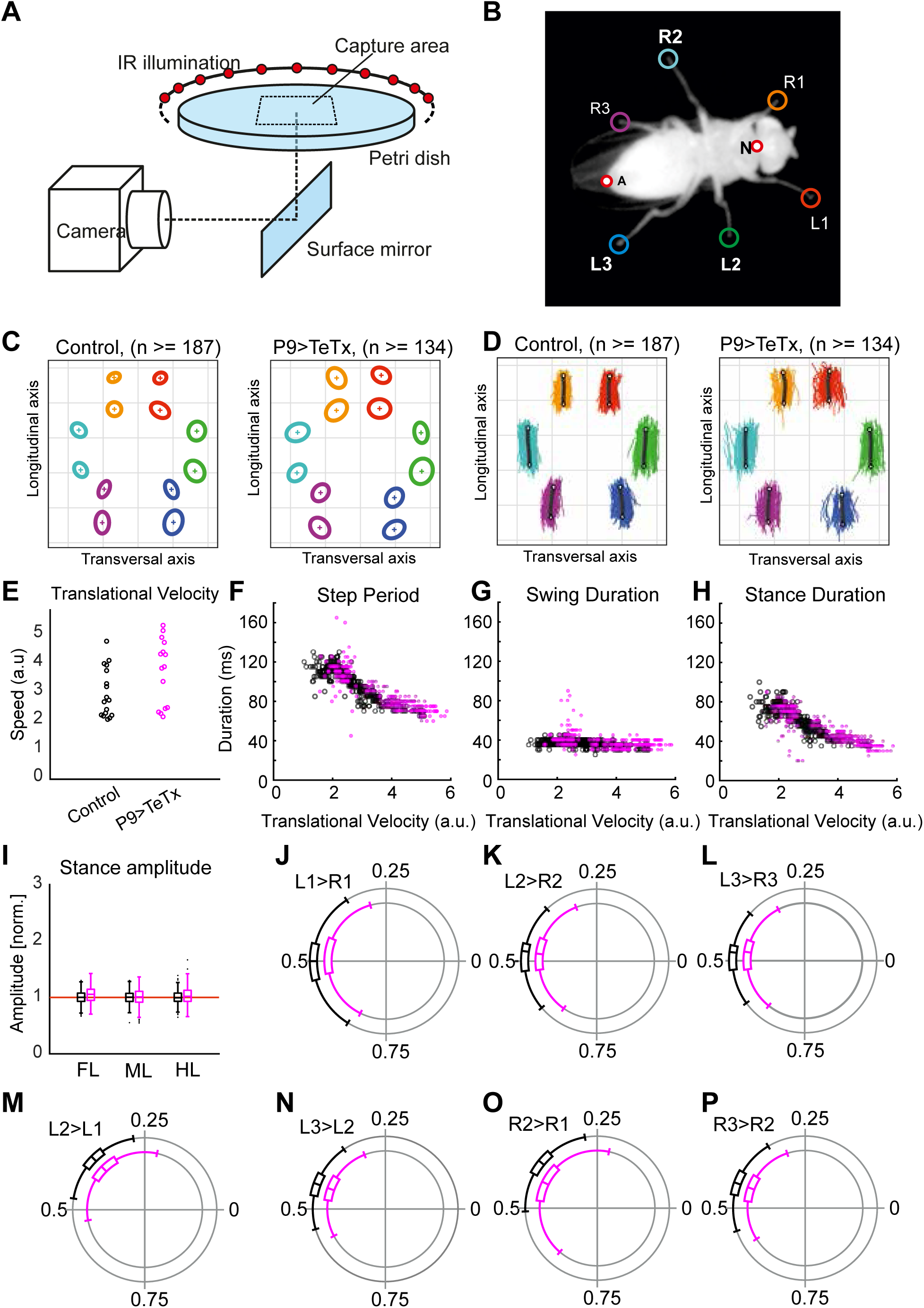
(A) High resolution walking analysis setup (B) Tracked frame showing leg positions tracked with corresponding labels. (C) Leg contact positions for each of the 6 legs of control (left) and P9>TeTx (right). (D) Stance trajectories for each of the 6 legs of control; n=134-187 steps from 4-5 flies per genotype (E) Translational Velocity of controls (black) and P9>TeTx (magenta) n = 15-17 trials. (F-H) Middle leg step period (F), swing duration (G) and stance duration (H) as a function of translational velocity of controls (black) and P9>TeTx (magenta). n=134-187 steps. (I) Stance amplitude of controls (black) and P9>TeTx (magenta). n=134-187 steps from 4-5 flies per genotype. (J-O) Phase difference plots for pairs of legs as per labels given in (B) showing individual steps as filled circles and mean values as vector line for (grey) controls and (magenta) P9>TeTx, n=134-187 steps from 4-5 flies per genotype.

**Figure S5 (related to Figure 5).**
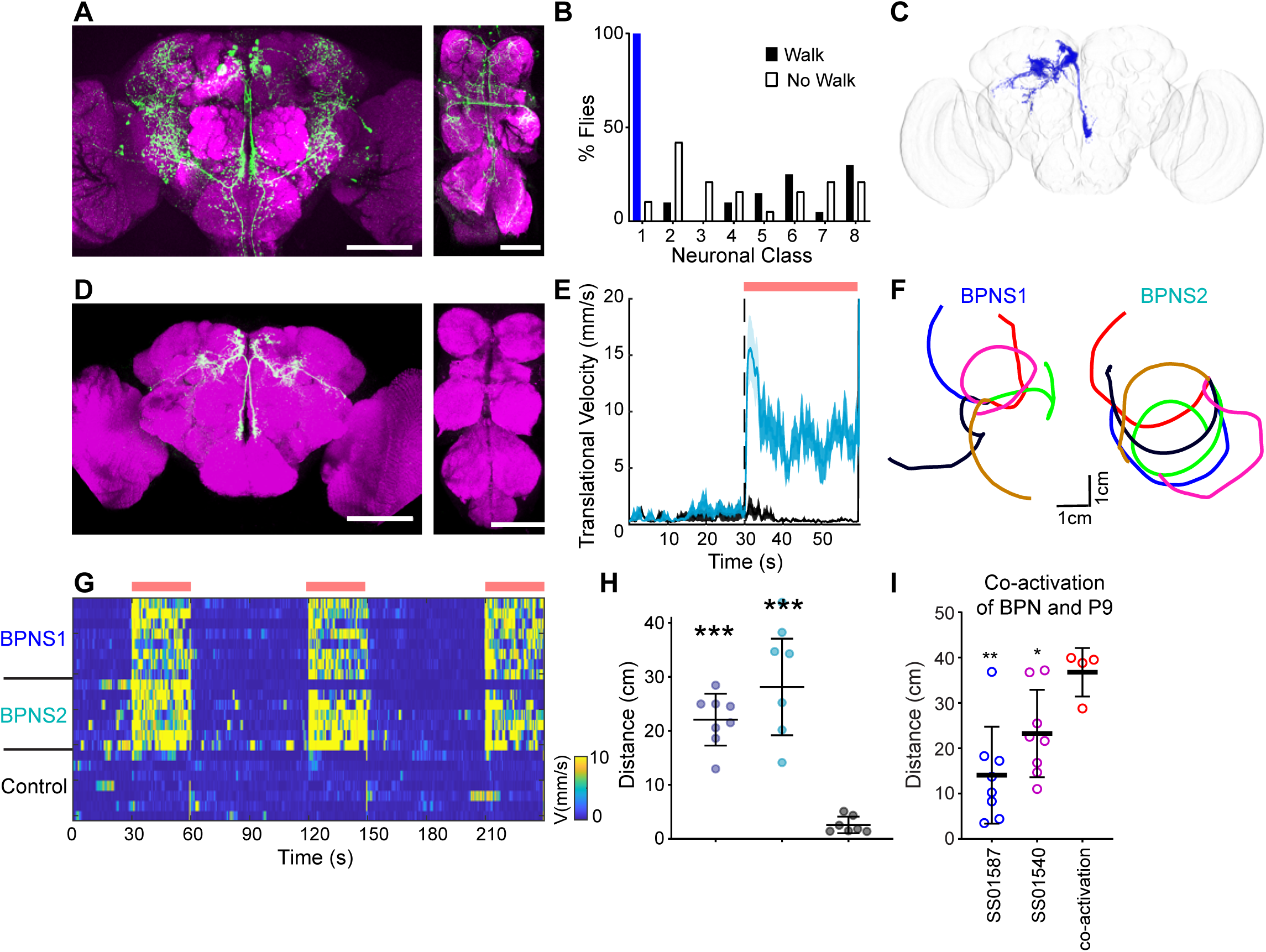
(A) SS01587>CsChrimson-mVenus expression in brain (green, left) and VNC (right). (B) Percentage of flies expressing CsChrimson-mVenus in 8 SS01587 cell-types for flies that initiated walking (filled bar) or did not (open bar) upon CsChrimson activation. n = 20 “Walk” and n=19 “No Walk”, Fisher’s exact test, ***p<0.001. (C) BPN segmented image. (D) BPNS2 split gal4 line expression data, CsChrimson-mVenus (green) and neuropil (magenta) for the central brain and VNC showing exclusive expression in BPNs. (E) Average trial translational velocity (mean ± SEM) in the grooming assay for BPNS2>CsChrimson flies (cyan) and controls (black, same experiment as in Figure 5B hence same controls. (F) Example walking trajectories showing 4 seconds after lightON for BPNS2, 7 flies. (G) Transient activation of walking in grooming flies on BPN activation using BPNS1 or BPNS2 depicted as velocity heatmaps for individual flies (red bars indicate light ON period). n=7-8 flies/genotype; controls are negative split combination). (H) Average distance during light ON period traveled by grooming flies while activating BPN splits compared to controls (n=7-8, ANOVA and Dunnett’s multiple comparison, ***p<0.001). (I) Example walking trajectories showing 4 seconds after lightON of BPN-S1, 7 flies. (J) Total distance traveled by grooming flies while co-activating SS01540 and SS01587 compared to individual activation (n=4-8, ** p<0.01, * p<0.05 Mann-Whitney Test).

**Figure S6 (related to figure 6).**
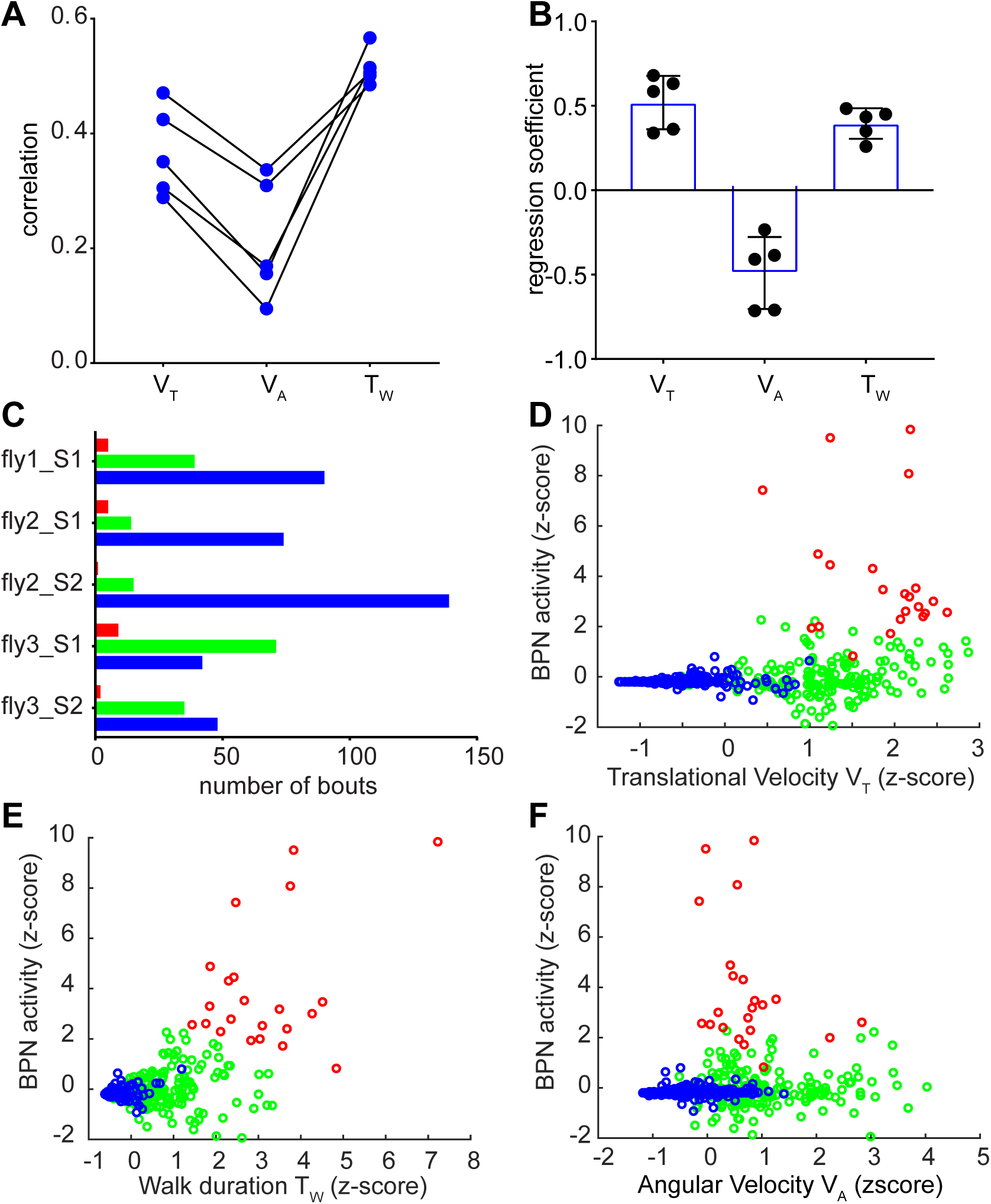
(A) Correlation coefficients for each BPN soma activity for example fly in Figure 6B-H, with regressors of translational velocity (V_T_), angular velocity (V_A_), walking bout duration (T_W_) and straightness index (SI). values for each soma can be followed by connecting lines. (B) Regression coefficients from linear regression model for example fly in Figure 6B-H (C) Distribution of different bout classes across 3 flies and 5 imaging sessions. Bouts color coded as in Figure 6 I,J (D-F) k-means clustering results from Figure 6I projected in 2D space of mean BPN activity versus (D) mean bout translational velocity, (E) bout walking duration and (F) mean bout angular velocity

**Figure S7 (related to Figure 7).**
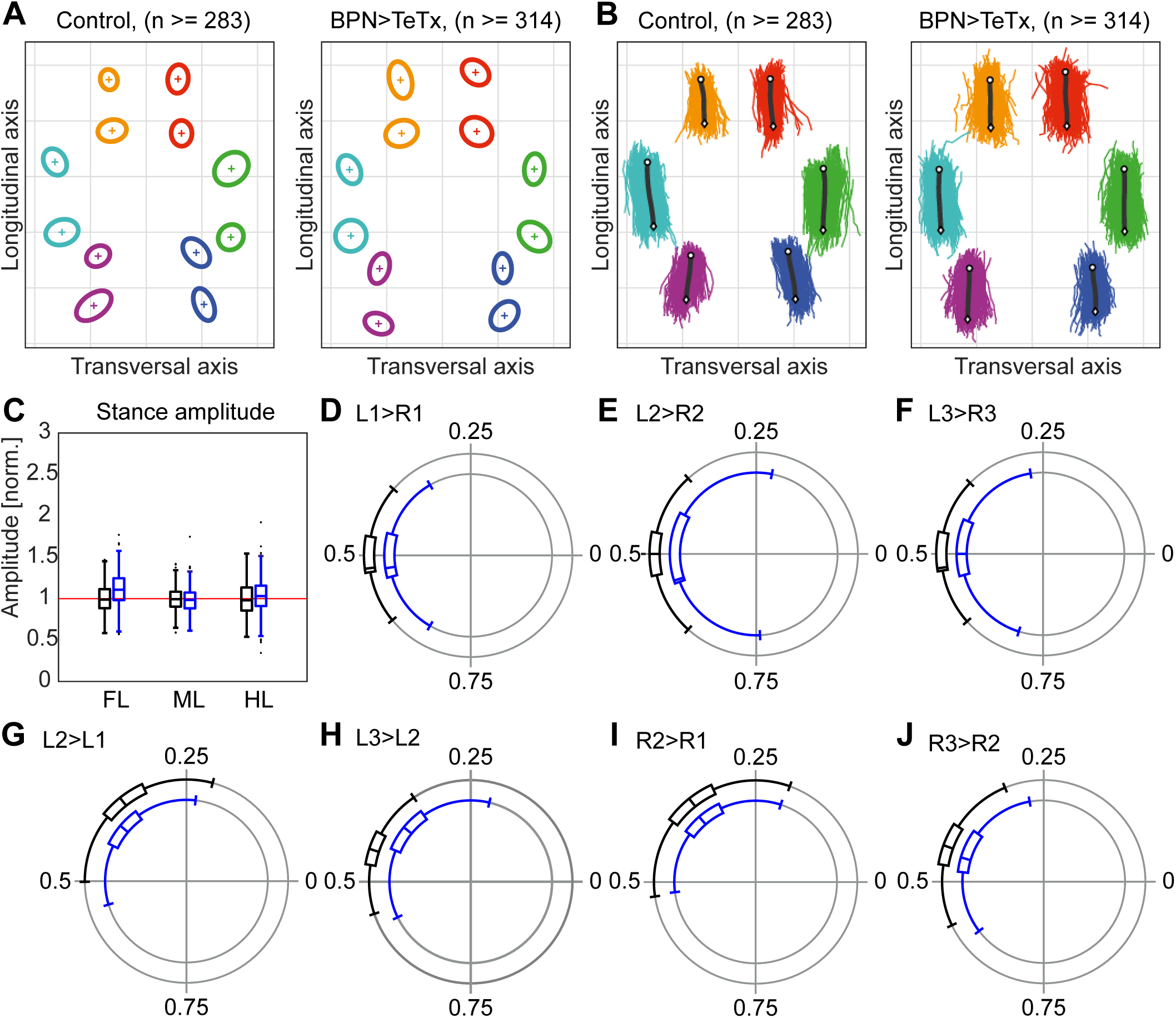
(A) Leg contact positions for each of the 6 legs of control (left) and BPN>TeTx (right). (B) Stance trajectories for each of the 6 legs of control and BPN>TeTx flies; n=134-187 steps (C) stance amplitude of (black) controls and (blue) P9>TeTx flies. (D) (J-O) Phase difference plots for pairs of legs as per labels given in (Figure S4B) showing individual steps as filled circles and mean values as vector line for (grey) controls and (blue) BPN>TeTx, n=134-187 steps.

## Supplementary Videos

**Video S1 (related to Figure 1):**

Transient activation in grooming flies for control (left), SS01540>CsChrimson (middle) and BPN-S1>CsChrimson (right) flies. Red dot at top left indicates light on (650nmLED)

**Video S2 (related to Figure 1):**

Transient activation in copulating flies for control (left), SS01540>CsChrimson (middle) and BPN-S1>CsChrimson (right) flies. Red dot at top left indicates light on (650nmLED)

**Video S3 (related to Figure 2):**

Transient activation of left P9 (left) or right P9 (right) using stochastic labeling of SS01540>stop>CsChrimson flies. Red dot on top left indicates light on (650nmLED)

**Video S4 (related to Figure 4):**

Control male (left) or P9>TeTx male (right) pursuing a remote-controlled (P9>CsChrimson) virgin female which is controlled using optogenetic activation indicated by red dot at top left. (650nmLED). Video speed is 2x real speed.

## STAR Methods

### LEAD CONTACT AND MATERIALS AVAILABILITY

Kristin Scott

### EXPERIMENTAL MODEL AND SUBJECT DETAILS

*Drosophila melanogaster* control and transgenic lines used in this study are described in detail in Key Resources table. Table S1 documents genotypes used for each figure. Flies were raised in vials containing standard cornmeal-agar medium supplemented with baker’s yeast and incubated at 25°C with 60% humidity and 12h light/dark cycle throughout development and adulthood unless otherwise stated. The behavioral screen and all optogenetic activation experiments were performed in female flies aged 7-10 days unless otherwise stated. For stochastic activation experiments, crosses were set at 19°C. and the larvae were heat shocked during development for 1.5 to 2 hrs at 37°C. All optogenetic activation experimental flies were collected on retinal food and again transferred to fresh retinal food 1-2 days prior to testing. Retinal food contains standard fly food with freshly added 400 μM all-trans retinal. These flies were kept in the dark from eclosion until test. The courtship experiments were performed using 8-10 day old test males paired with 5-6 day old virgin females. The males were collected as virgins and raised in isolation in 1.5 ml eppendorf vials with ~ 0.75ml standard food and holes for air exchange.

### METHOD DETAILS

#### Fly Genetics

All DN split-Gal4 lines used in the screen were generously provided pre-publication by S.Namiki and G.Card. The VT and GMR generation 1 Gal4 lines lines were obtained from Vienna Drosophila Resource Center (VDRC) and Bloomington Drosophila Stock Center (BDSC) respectively. The new split-Gal4 lines for P9 and BPNs were created using color MIP masked search on the entire registered image database (Dionne et al., 2018; Otsuna et al.2018; Tirian and Dickson, 2017) made available by Janelia Research Campus. Candidate drivers targeting the neurons of interest were identified and corresponding split-Gal4 hemi-drivers were obtained from BDSC. Sources for other reagents used are described in Table S1 and Key resources table.

#### Activation Screen

All behavioral experiments were performed in a temperature and humidity-controlled room (25°C, 60% humidity). The behavioral arenas were designed by pouring 1.5% agarose gel in a regular 150 mm petri-plate with 3D printed acrylic molds designed for generating bowl-shaped behavioral arenas. The arenas were covered with a glass ceiling painted with Sigmacote. Four bowl-shaped arenas of 44 mm diameter were imaged with one camera (JVC camcorder GZ-E300AU) at a resolution of 1920×1080 and 30 fps. Four cameras operated in parallel during the screen. The arena was backlit with custom designed bright white LED panels and the light was on throughout the assay. Intensity was adjusted using MDN>CsChrimson (Bidaye et al., 2014) as a positive control. For long duration continuous stimulation, white light of 0.5 mW//cm^2^ at the arena walking surface was sufficient to observe CsChrimson phenotypes. Powdered and clean flies were tested for each genotype. The powdered flies were coated with fine yellow dust (Reactive Yellow 86, Cat # sc-296260, (Seeds et al., 2014) just prior to loading them into the behavioral arena. A 7 minute video was acquired after flies acclimatized to the behavioral arena for 3 minutes (dim lights).

#### Transient Activation Setup

A custom LED plate was designed as a backlight for optogenetic experiments. The plate (150 mm X 150 mm) consists of high density high power LEDs (10×10 LED positions). 3 LEDs per position, 870nm, 630nm and 530nm, each with an independent intensity control, a switch input and a TTL input for pulsing at desired frequencies. The panel was placed underneath a diffuser sheet to provide even illumination at the arena walking surface. The 870nm IR LED was switched ON at low intensity throughout the assay. For CsChrimson activation experiments, 630nm LED was adjusted to output ~4.5 mW/cm^2^ intensity at the arena walking surface and was transiently switched on and pulsed at 50Hz (5ms pulse width) via the TTL input unless otherwise stated. For the green light walking induction experiments, the 530 nm green LED was adjusted to output ~1.8mW/cm^2^ intensity at the arena walking surface and transiently switched on during the assay. Videos in this setup were acquired using a FLIR Blackfly S camera (FL3-U3-13Y3M-C) at a resolution of 1280 × 1024 at 30 fps. The camera was fitted with an adjustable focus lens (LMVZ990-IR) and NIR bandpass filter (Midopt BP850) to allow IR imaging without artifacts from visible light. All transient activation experiments (unless otherwise stated) were performed in the dark.

#### Transient Activation Grooming Assay

Powdered flies loaded in bowl-shaped arenas described above were assayed for walking initiation on transient activation in the above setup. The light stimulation protocol consisted of 60s OFF 30s ON sequence, repeated 3 times. For analysis, 30s OFF followed by 30s ON was considered as one trial.

#### Transient Activation Copulation Assay

Virgin test females were paired with Canton S wild type males in the bowl-shaped arena described above and allowed to begin copulation. Once copulation was initiated, the light stimulation protocol of 30s OFF 10s ON was repeated 3 times. Longer assay duration or light ON periods often led to unmounting. For analysis, 8s OFF followed by 8s ON was considered as one trial. In the event of rare unmounting during the assay, that video was discarded from the analysis.

#### Courtship Assay

We paired Canton S virgin females with test males in bowl-shaped arenas described above placed on top of white LED backlight (0.5 mW//cm^2^). 10 min videos were acquired using JVC camcorders (GZ-E300AU) at 1920x 1080 resolution and 30 fps.

#### Remote-controlled female courtship assay

Virgin SS01540>CsChrimson(attp18) females (aka remote-controlled females or P9>Chr) were paired with test males in bowl shaped arenas and placed in the transient activation setup. Diffuse room lighting provided enough light to increase courtship compared to lights OFF, indicating that visual aspects of courtship were functional. During the assay, the remote-controlled females were triggered to initiate high forward and angular velocity walking by transiently switching on the 630nm light (10s OFF 20s ON for a total of 10 minutes). 10 minute videos were recorded using the FLIR BlackFlyS camera at 1280 × 1024 and 30 fps.

#### Flat arena Courtship Assay

Canton S virgin females were paired with test males in flat floor arenas designed to increase complex following events. These arenas are 50mm diameter and 3 mm high with serrated walls. Arena floor is a diffuser sheet placed on hard clear acrylic sheet. The arenas were backlit with white LEDs (0.5mW/cm^2^) and 10 min videos were recorded using JVC camcorder at 1920x 1080 resolution and 30 fps.

#### Freezing Assay

For reproducing the P9 freezing phenotype we designed an arena as similar to the published one (Zacarias et al., 2018) as possible. The main difference from our usual bowl shaped arena was that the floor of these arenas was hard flat acrylic instead of curved agarose floor. These arenas were identical in dimensions to the flat shaped courtship arena described above except these arenas did not have serrated walls, so that flies could also walk on the arena walls, similar to Zacarias et al assay. The LED intensity and pulsing durations were adjusted to be similar to Zacarias et al assay, viz. ~2.5 mW/cm^2^ intensity and 10 repetitions of 2s LED on (continuous on without pulsing) followed by 10s LED off period.

#### Green Light Induced Walking Assay

Test flies were aspirated in flat arenas and placed in the transient activation setup. The flat courtship arenas were used, so that baseline walking velocity was low allowing us to see an increase in walking velocity with green light. Green light was switched on transiently (60s OFF 30s ON) for three trials and videos were recorded on FLIR BlackFlyS camera at 1280 × 1024 at 30 fps.

#### BPN walking assay

Same conditions as green light assay, except that green light was not turned on during the 10 minute video recording.

#### High resolution walking assay

A schematic of the free-walking setup is shown in Figure S4A. It consisted of an inverted glass petri dish that we used as a transparent arena (diameter 80 mm) held by a circular frame with a cutout below the dish. The cutout provided an unobstructed bottom view of the arena. A surface mirror was placed below the arena at a 45° angle. Using this mirror and an infrared (IR)-sensitive high-speed camera (model VC-2MC-M340; Vieworks, Anyang, Republic of Korea), we captured a bottom view of a central quadratic area on the surface of the arena of approximately 30 x 30 mm, with a resolution of 1000 by 1000 pixels, a frame rate of 200 Hz, and an exposure time of 200 µs. Illumination was provided by a ring of 60 IR light-emitting diodes (LEDs, wavelength: 870 nm) arranged concentrically around the arena and emitting their light mainly parallel to the arena’s surface. This resulted in a strong contrast between background and fly (see Fig. S4B). Contrast and homogeneity were further increased by equipping the camera’s lens with an IR-pass filter (cut-off frequency: 760 nm) that blocked all ambient visible light. The LED activity was pulsed and synchronized to frame acquisition of the camera. To prevent flies from escaping, the arena was covered with a watch glass that established a dome-shaped enclosure, similar to an inverted FlyBowl (Simon & Dickinson, 2010). To prevent flies from walking upside down on the watch glass, we covered its inside surface with SigmaCote (Sigma-Aldrich, St. Louis, MO). Prior to an experiment, a single fly was extracted with a suction tube from its rearing vial, cold-anesthetized for approximately one minute, and placed onto the arena, which was then covered with the watch glass. Flies were allowed to regain mobility and then acclimatize for ~15 minutes, after which video acquisition was started.

Flies walked spontaneously for several hours in the arena and frequently crossed the capture area. Video data of this area was continuously recorded into a frame buffer of 5-10s duration. During an experiment, custom-written software functions evaluated the recorded frames online and determined if a fly was present at a particular time and if it had produced a continuous walking track that had a minimum length of 7 BL (body lengths) and a minimum walking speed of 2 BL s-1. Once the fly had produced such a track and then either stopped or left the capture area, the contents of the frame buffer were committed to storage as a trial for further evaluation and analysis. Video acquisition and online evaluation during experiments were implemented in MATLAB 2018b (The MathWorks, Natick, MA).

#### Immunohistochemistry

Staining of fly brain and ventral nerve cord was performed as described previously (Yu et al., 2010). Briefly, flies were dissected in cold PBS and brains and VNCs were fixed in 4% PFA. This was followed by PBST washes, then blocking in normal goat serum, followed by primary antibody incubation for 2 days, PBST washes and secondary antibody incubation for another 2 days. The stained tissues were then mounted in Vectashield and imaged under LSM 780 system using the 1P excitation and emission corresponding to the secondary antibodies used. Antibodies are as described in Key resources table. Primary antibodies were used at 1:500 or 1:1000 dilution and secondaries were used at 1:500 dilution.

#### Neuron morphology segmentation

For P9 and LC9 neuron segmentation we used the publicly available split-gal4 resource (Namiki et al., 2018; Wu et al., 2016) and registered it on the JFRC2 template using the CMTK toolkit. For pC1 we used the JFRC2 template registered pMP-e image obtained from virtualFlyBrain resource (Cachero et al., 2010; Osumi-Sutherland et al., 2014). For BPN, we registered our nc82 co-stained image of BPN-S1>CsChrimson on the JFRC2 template using the CMTK toolkit (Jefferis et al., 2007). The neuron segmentations were created using the registered images in VVD software (Otsuna et al., 2018). VVD was also used for visualizing potential overlap between the segmented neurons and creating the segmented neuron figures.

#### CsChrimson quantification

8-9 day old female flies were cold anesthetized and brains were dissected in cold Artificial Hemolymph (AHL) solution (Wang et al., 2004). The brains were then transferred on a PLL coated coverslip posterior side up and immersed in AHL. CsChrimson-mVenus expression was detected using 514nm confocal laser scanning of region of interest under a 20x NA 0.8 objective lens on a LSM 880 Zeiss microscope. For P9 CsChrimson analysis, zoomed in 3D stack comprising the entire soma was captured. For gamma lobe CsChrimson analysis, 3D ROI consisting of the neurites was imaged. This entire data set (Figure S2 G) was obtained by imaging age matched flies using identical imaging parameters during the same imaging session.

#### Functional Connectivity

Functional connectivity experiments were performed in freshly dissected explant brains under a LSM 880 two photon microscope using 20x NA =0.8 objective lens. Brains and connected VNC were dissected and imaged in extracellular saline (ECS) solution (103mM NaCl, 3mM KCl, 5mM N-tris(hydroxymethyl) methyl-2-aminoethane-sulfonic acid, 10mM trehalose, 10mM glucose, 2mM sucrose, 26mM NaHCO3, 1mM NaH2PO4, 1.5mM CaCl2, and 4mM MgCl2, adjusted to 275 mOsm, pH equilibrated near 7.3 when bubbled with carbogen). The explant was placed on PLL coated coverslip in an imaging chamber (ALAMS-518SWPW). ECS bubbled with carbogen was perfused over the brains throughout the imaging session. To activate Chrimson, 650 nm red light was delivered to the sample through the objective. A BP excitation filter centered around 655nm and custom notch dichroic (Chroma ZT656dcrb) delivered the red light on the sample and the dichroic at the same time allowed transmission of the 920nm imaging laser as well as the green GCaMP signal back to the detectors. The activation light LED was controlled via TTL inputs synchronized to the imaging session using ZEN2.0 software and an external trigger box plus a signal generator. A given session consisted of three to five activation trials consisting of 30s OFF 10s ON periods. During the ON period, the LED was pulsed at 50 Hz (2ms pulse duration). 512 x 512 pixel images were acquired at 3 to 4 Hz. A high intensity two color 1P confocal stack was acquired at the end of the imaging session to confirm expression. In case of samples that showed GCaMP responses, the brains from the same batch of flies were immunostained to confirm there was no co-expression of Chrimson and GCaMP in the same neuron.

#### In-vivo Imaging

Imaging in tethered walking flies was performed on a 3i spinning disc confocal system with a 20x water immersion objective (NA 1.0). Male flies (age 5-7) were cold anesthetized and attached on a custom holder, modified version of (Weir et al., 2016). The head was positioned such that the ocelli were on the top and covered with AHL. The holder ensured that only the superior dorsal part of the head was immersed in the solution leaving the antennae and proboscis dry. The proboscis was glued to reduce brain movement. A small hole was created in the cuticle encircling the ocelli region and air sacs were removed. The holder was then placed under a 20x objective and imaged using a 488nm laser. This dissection revealed BPN soma; however, since the soma are positioned along the curved top surface of the brain, not all soma could be imaged in all samples. Once the imaging ROI was centered, an air supported ball was positioned under the fly using micromanipulators. The track-ball (polypropylene, diameter: 6 mm, Spherotech GmbH, Germany). freely floated on a controlled air stream through a ball holder (Berendes et al., 2016; Seelig et al., 2010). Two cameras placed at right angle to each other were used to carefully align a square sleeve placed on the ball holder such that it aligned with the axis of the fly (Green et al., 2017). This was later used to determine the coordinate frame of the fly via FicTrac (Moore et al., 2014). The ball was illuminated with 850nm IR LED and imaged using a FLIR BlackFly S Camera and lens mentioned in the above section. Video was acquired at 400 x 400 resolution and 50fps and was synchronized to the imaging session via slidebook software and a TTL trigger input to the camera. Once the fly was acclimatized to the ball and started walking, a volume consisting of the BPN soma was imaged at frame rate of 0.5-0.9 Hz. Imaging sessions of walking flies usually lasted 5-7 minutes. The synchronized GCaMP imaging and ball tracking data was then analysed offline.

### QUANTIFICATION AND STATISTICAL ANALYSIS

All data analysis was performed in Matlab and statistical tests were performed in Graphpad Prism. Graphs were plotted either in Matlab or Graphpad.

#### Analysis of walking during activation assays

Videos were processed using a custom batch processing script that cropped videos into single arena videos and uncompressed to avi format. These preprocessed videos were then tracked using FlyTracker software (Eyjolfsdottir et al., 2014). The tracker returned the centroid position of the fly, orientation and instantaneous translational velocity. For the activation screen, we only used the translational velocity parameter. Since the screen spanned several days of experiments, we recorded a control video (UAS-CsChrimson x w1118) every day of both clean and powdered flies (n = 16 each). All test fly data was then normalized with respect to the day control for visualizing outlying clusters (Figure 1B and S1A). We used the orientation attribute to determine the signed or absolute angular velocity of the fly. All plots except for stochastic activation experiments report the absolute angular velocity of the fly. We calculated angular velocity, translational velocity, distance traveled and total angular rotation (area under the curve of absolute angular velocity) for each trial and then averaged the trials for plotting and statistical analysis of per fly attributes. Distance and total rotation values were primarily used for statistical analysis and velocity profiles were used for showing the dynamics of the activation phenotype. In all transient activation data figures, n represents number of flies and the values for n and statistical analysis is described in the figure legends.

#### Courtship Assays

The courtship videos were preprocessed and tracked using FlyTracker as described above. The two-fly tracking output was manually checked for fly identity mismatch using “visualizer” script in the FlyTracker software and any mismatch was corrected. The centroid positions and orientations of the flies were used to calculate the distance between the two flies and the angle between the male and female as per previous reports (Ribeiro et al., 2018). The following Index was defined as fraction of pre-copulation time spent by the male within 5 mm of female and oriented towards the female (angle < 30) (Ribeiro et al., 2018). Velocity correlation was the correlation between the translational velocities of male and female fly in a given courting pair before copulation. Probability density functions of distance and angle between the courting pair were calculated per courting pair and then plotted as mean ± SEM for visualization.

#### CsChrimson expression quantification

Mean fluorescence intensity was quantified for each ROI (P9 soma or gamma lobe neurites). Data for all flies imaged was reported. The gamma lobe expression was only present and quantified in the SS01540 line.

#### Functional Connectivity

Mean ROI fluorescence of P9 soma and a background ROI was calculated using ImageJ. Background subtracted fluorescence intensity was used to calculate ΔF/F0 where F0 is calculated as mean fluorescence during 10s prestimulation light OFF period. We plotted data for 10s OFF 10s ON and 10s OFF for each trial. Area under the curve during light ON period was used for statistical analysis.

#### In-vivo imaging

GCaMP6 imaging data was analyzed using open source and custom Matlab scripts. The NormCorre Matlab package (Pnevmatikakis and Giovannucci, 2017) was used for image registration to remove movement artifacts. ROIs were manually drawn around each visible BPN soma, ignoring somas close to Z-stack boundary plane to avoid movement artifact, and mean ROI fluorescence was extracted. This pre ROI fluorescence data was then bleach corrected using the bleach correction script from CaImAn-MATLAB toolbox (Giovannucci et al., 2019). The ΔF/F0 values (threshold 0.1 and 10-90 percentile range > 0.2) for each ROI were used to filter out soma that did not show activity during the imaging session. 5 of 9 soma for fly1, 3 of 5 soma for fly2 and 6 of 10 soma for fly3 we defined as active. The bleach corrected BPN soma fluorescence data was then z-scored and used for further analysis. The mean BPN activity data was mean of z-scored bleach corrected fluorescence for all active soma.

The ball movement was tracked offline using FicTrac software (Moore et al., 2014). FicTrac directly provided the position of the fly in a virtual space as well as the instantaneous translational velocity and virtual heading of the fly. These attributes were used for calculating angular velocity, walking bout duration and straightness index. Straightness index (SI) was a modified version of previously reported definition (Cruz et al.). In order to calculate straightness without any effect of velocity we defined SI as a pure spatial parameter. For every point on the trajectory, 12mm trajectory segment centered around that point was extracted (spatial window) and SI for that segment was calculated as trajectory length divided by the sum of deviations from an ideal path (i.e. a straight line joining the first and last points in that window). Straight paths have small deviations from ideal path and hence larger value of SI. Also because SI was not defined using a temporal window, we did not include SI as an attribute in the linear model or clustering of other time series attributes.

The low frame rate GCaMP imaging data were upsampled (interpolated to high frame rate of 1kHz and then digitized to 50 Hz) to match the ball tracking frame rate for data analysis. The behavioral attributes were convolved using a GCaMP6s kernel (Turner-Evans et al., 2017) and the obtained regressors were used for calculating correlations and generating a linear regression model. Although correlations of BPN activity with behavioral regressors could be obtained, the slow <1Hz imaging frame rate made it difficult to make any claim about the lag between the two variables.

For pooled data analysis across flies and imaging sessions, data for each imaging session was first divided into walking bouts defined as any continuous walking event (V_T_ > 2 mm/s). Then the z-scored mean BPN activity and behavioral attributes (V_T_, V_A_, T_W_) were clustered using k-means clustering into 3 clusters. To visualize the BPN activity dynamics for each cluster, we aligned the BPN activity data to walking bout initiation frame and plotted mean BPN activity before and after bout initiation for each cluster separately as mean ± SEM bounded plots (Figure 6J).

#### High resolution walking analysis

Prior to data analysis, we selected walking sequences which had low within-trial variability in walking speed (translational velocity), in which flies walked in a straight line, and which contained at least five consecutive steps. These pre-selected sequences served as the basis for further analysis of low-level walking-related parameters. First, the position of the fly throughout a sequence was determined automatically. In brief, each video frame was converted into a binary image (black background, bright fly), in which, following a simple threshold operation, the fly was detected as the largest bright area. The walking speed was calculated as changes of the center of mass of this area over time. We then used this positional information to crop the fly from the original 1000 by 1000-pixel video. These smaller and fly-centered video sequences were used for the annotation of eight different body parts in every video frame: the tarsal tips of all six legs, the neck, and the posterior tip of the abdomen (Fig. S4B). This step of the annotation was done automatically in DeepLabCut (DLC, Mathis et al, 2018); to use this approach we trained and evaluated DLC with a data set of 1000 manually annotated video frames (10 flies, 100 exemplary frames each), that were similar to the ones we recorded during the experiments described here. One half (500 frames, 10 flies, 50 frames each) of this set was used for training DLC, the other half was used to evaluate its performance. Performance of DLC was generally very good; however, to ensure high-quality annotations we inspected the results visually and, if necessary, corrected rare mis-annotations manually.

To determine the times of lift-off and touch-down for each leg in a walking sequence, the DLC-determined positions of the tarsal tips were transformed into a world-centered coordinate system. In this coordinate system, a leg tip is stationary, i.e. has a speed of zero, when the leg is touched down (here, defined as the stance movement) and moves markedly with regard to the ground when it is lifted off (here, defined as the swing movement). These measures and an empirically determined threshold were used to distinguish between swing and stance movement. Transitions between these two were defined as touch down and lift off events, respectively, and the positions of the tarsal tip at these times were defined as the anterior and posterior extreme positions (AEPs and PEPs) in fly-centered coordinates, respectively. A single step of a particular leg was then defined as its movement between two subsequent PEPs; its period was defined as the time difference between two subsequent PEPs. Swing movement and duration were defined as the movement and the time difference, respectively, between a PEP and the subsequent AEP; stance movement and duration were defined as the movement and the time difference, respectively, between an AEP and the subsequent PEP. A stance trajectory was defined as the complete path of a tarsal tip in fly-centered coordinates between an AEP and the subsequent PEP. Step amplitude was defined as the distance between a PEP and the subsequent AEP. Stance linearity was calculated as the root mean squared error (RMSE) between an actual stance trajectory and a straight line between this stance trajectory’s AEP and PEP; the higher this measure is the stronger the deviation from a straight line. Average AEPs and PEPs were defined as the arithmetic mean of all AEP and PEP position vectors, respectively; the standard deviation of these positions were estimated as a bivariate distribution. Stance trajectories were averaged by first re-sampling all n trajectories to 100 equidistant positions and then calculating the arithmetic mean for each set of n-by-100 data points. To facilitate comparison between control and experimental condition we normalized all individual step periods, swing durations, stance durations, stance amplitudes, and stance linearity values to the arithmetic mean of the control condition.

Phase relationships, i.e. phase differences, were calculated for all ipsilaterally or contralaterally adjacent leg pairs. This resulted in seven phase relationships: three contralateral leg pairs (front, middle, and hind legs), as well as four ipsilateral leg pairs (hind and middle legs, and middle and front legs, respectively). For each leg pair a reference leg was selected. For contralateral phase relationships this was a left leg, for ipsilateral phase relationships this was the posterior leg. For each complete step of the reference leg (i.e. PEP to PEP) we calculated its instantaneous phase as a value that linearly increased from 0 to 1 during the step. We then calculated the phase relationship between the two legs as the phase value at the times of PEPs in the non-reference leg; this is equivalent to the phase difference between these legs. A value of 0 indicates synchronous lift off, for instance, while a value of 0.5 would indicate exactly anti-phase. The phase between to legs is indicated by, for instance, R3>R2, where the right hind leg (R3) is the reference leg and R2 (the right middle leg) is the non-reference leg.

All annotations and calculations, apart from DLC-based functions, were carried out with custom-written functions in MATLAB 2018b.

### DATA AND CODE AVAILABILITY

Raw and processed data along with custom Matlab analysis scripts will be made available upon request. Design files for custom parts will be available upon request.

